# Ana1 recruits PLK1 to mother centrioles to promote mitotic PCM assembly and centriole elongation

**DOI:** 10.1101/2020.08.11.244194

**Authors:** Ines Alvarez-Rodrigo, Alan Wainman, Jordan W. Raff

**Affiliations:** The Sir William Dunn School of Pathology, University of Oxford, South Parks Road, Oxford OX1 3RE

**Keywords:** PLK1, Polo, Centrosome, Centriole, PCM, Ana1, Cep295, mitosis

## Abstract

Polo kinase (PLK1) is a master cell cycle regulator that is recruited to various subcellular structures by its Polo-Box domain (PBD), which binds to phosphorylated S-pS/pT motifs. Polo has multiple functions at centrioles and centrosomes, and we previously showed that phosphorylated Sas-4 initiates Polo recruitment to newly formed centrioles, while phosphorylated Spd-2 recruits Polo to the mitotic Pericentriolar Material (PCM) that assembles around mother centrioles. Here, we investigate whether additional proteins recruit Polo to centrioles and/or centrosomes, and find that Ana1 (Cep295 in mammals) helps recruit Polo to mother centrioles. If this function is impaired, mother centrioles can still duplicate and disengage from their daughters, but they can no longer efficiently assemble a mitotic PCM or elongate their centrioles in G2. Thus, Ana1 is part of a sequential phosphorylation cascade that recruits Polo to centrioles to drive mitotic centrosome assembly and centriole elongation in G2, but not centriole duplication or disengagement.

## Introduction

Polo/PLK1 kinase is an important cell cycle regulator (Pintard and Archambault, 2018). During mitosis, it is recruited to several different locations in the cell—most prominently to centrosomes, kinetochores and the cytokinesis apparatus—where it performs multiple functions (Colicino and Hehnly). PLK1 is commonly recruited to these locations by its Polo-Box-Domain (PBD) (Lee et al., 1998; Liu et al., 2004; Reynolds and Ohkura, 2003), which binds to phosphorylated S-pS/pT motifs in target proteins (Elia et al., 2003a; b). Mutating the first Ser in the PBD-binding motif to Thr strongly reduces PBD-binding in vitro and also in vivo (Elia et al., 2003a; b; Decker et al., 2011; Joukov et al., 2014; Meng et al., 2015).

PLK1 has several key roles at centrosomes. These organelles are important microtubule (MT) organising centres, and are formed when a pair of centrioles (comprising an older mother and younger daughter) recruit a matrix of pericentriolar material (PCM) around the mother centriole (Conduit et al., 2015b). Centrosomes help to organise the poles of the mitotic spindle, and towards the end of mitosis the mother and daughter centrioles disengage from each other, so that each new cell inherits two closely-spaced, but disengaged centrioles. PLK1 is essential for disengagement (Tsou et al., 2009; Kim et al., 2015), and also for the subsequent maturation of the daughter centriole into a new mother centriole that is itself capable of duplicating and organising PCM (Loncarek et al., 2010; Wang et al., 2011; Kong et al., 2014; Shukla et al., 2015; Novak et al., 2016). The old mother (OM) and new mother (NM) centrioles then both duplicate during S-phase by nucleating the assembly of a daughter centriole on their side. It seems that PLK1 is not essential for centriole duplication *per se*, which involves the assembly of a central cartwheel structure on the side of the mother that then recruits centriole MTs and which is largely driven by the related kinase Plk4 (Breslow and Holland, 2019; Gönczy and Hatzopoulos, 2019), but it is required for the subsequent growth of the centriole MTs that occurs during G2 (Kong et al., 2020). After duplication in S-phase, the two centrosomes (each now comprising a duplicated centriole pair) are held together by a linker, and PLK1 plays an important part in linker disassembly and in promoting centrosome separation as cells prepare to enter mitosis (Bertran et al., 2011; Mardin et al., 2011; Smith et al., 2011).

During interphase, centrosomes organise relatively little PCM but as cells prepare to enter mitosis the PCM expands dramatically in a process termed centrosome maturation (Palazzo et al., 1999). PLK1 is recruited to the PCM and is an essential driver of this process (Lane and Nigg, 1996; Haren et al., 2009). Several PCM proteins involved in centrosome maturation have been identified as PLK1 targets. In vertebrate cells, PLK1 phosphorylates Pericentrin, which cooperates with Cdk5Rap2 to promote mitotic PCM assembly (Lee and Rhee, 2011; Kim and Rhee, 2014), whereas in flies and worms PLK1 directly phosphorylates Cnn or SPD-5 (functional homologues of Cdk5Rap2), respectively, which allows these proteins to assemble into a PCM-scaffold around the mother centriole that recruits other PCM proteins (Conduit et al., 2014a; b; Woodruff et al., 2015, 2017; Wueseke et al., 2016; Feng et al., 2017).

These studies indicate that PLK1 has at least five important functions at centrosomes: (1) It promotes centriole disengagement; (2) It promotes the maturation of the daughter centriole into a mother; (3) It promotes the growth of the daughter centriole in G2; (4) It promotes centrosome separation; (5) It promotes centrosome maturation. How PLK1 is recruited to centrosomes to execute these multiple functions is largely unclear, although some progress has recently been made in understanding how Polo is recruited to drive centrosome maturation. In vertebrate systems, Cep192 is required for efficient centrosome maturation (Gomez-Ferreria et al., 2007; Zhu et al., 2008) and is phosphorylated by Aurora A to create PBD-binding sites that recruit PLK1; this promotes the mutual activation of both kinases to promote mitotic PCM assembly (Joukov et al., 2010, 2014). In flies and worms, Spd-2/SPD-2 (the fly/worm homologue of Cep192) is concentrated at centrioles and centrosomes and its phosphorylation also helps recruit PLK1 to the mitotic PCM so that it can phosphorylate Cnn/SPD-5 (Decker et al., 2011; Alvarez-Rodrigo et al., 2019). In fly embryos, Spd-2, Polo and Cnn have been proposed to form a positive feedback loop that drives the expansion of the mitotic PCM around the mother centriole (Conduit et al., 2014b; Alvarez-Rodrigo et al., 2019). In this scenario, Spd-2 is initially recruited to centrioles and is phosphorylated during mitosis, allowing the Spd-2 to form a scaffold that can recruit other PCM proteins and that fluxes outwards from the mother centriole. The Spd-2 scaffold appears to be weak, and so it quickly dissipates; but if it can recruit Polo and Cnn, the Polo can phosphorylate the Cnn to form a Cnn-scaffold that both recruits PCM components and also strengthens the Spd-2 scaffold. This allows more Spd-2 to accumulate around the centriole, which in turn drives the recruitment of more Polo and Cnn—so forming a positive feedback loop. Thus, Cep192/Spd-2 proteins recruit PLK1/Polo to the PCM to drive centrosome maturation.

If fly Spd-2 is mutated so that it can no longer efficiently recruit Polo—by mutating all S-S/T motifs to T-S/T—Spd-2 can no longer recruit Polo to the PCM, but Polo is still recruited to the mother centriole, indicating that other proteins must recruit Polo to this location (Alvarez-Rodrigo et al., 2019). We previously showed that the centriole protein Sas-4 is phosphorylated by Cdk1 during mitosis on Thr-200, creating a PBD-binding site that recruits Polo to newly formed centrioles for the first time (Novak et al., 2016). This phosphorylation enables the new mother centrioles to recruit Asl (Cep152 in vertebrates), which in turn allows mother centrioles to both duplicate (as Asl helps recruit Plk4 to centrioles (Cizmecioglu et al., 2010; Dzhindzhev et al., 2010; Hatch et al., 2010; Kim et al., 2013; Novak et al., 2014; Shimanovskaya et al., 2014)) and to organise PCM (as Asl helps recruit Spd-2 to centrioles (Conduit et al., 2014b)). While it is possible that the single PBD-binding site in Sas-4 is sufficient to recruit Polo to mother centrioles, we suspected that other proteins must help to recruit the large amounts of Polo that are normally concentrated there. Here, we attempt to identify such proteins by mutating all the S-S/T motifs to T-S/T in several candidate proteins. We find that the centriole protein Ana1 (Cep295 in humans) normally helps recruit Polo to mother centrioles. Ana1/Cep295 is required for centriole maturation (Izquierdo et al., 2014; Fu et al., 2016; Tsuchiya et al., 2016), and in flies Ana1 helps recruit and/or maintain Asl at new mother centrioles (Fu et al., 2016; Saurya et al., 2016), so the centrioles in flies lacking Ana1 cannot duplicate or recruit mitotic PCM (Blachon et al., 2009; Fu et al., 2016; Saurya et al., 2016). Surprisingly, centrioles in which Ana1 cannot recruit Polo can still recruit Asl, can still duplicate and disengage, but they cannot efficiently recruit mitotic PCM or elongate during G2.

## Results

### Mutation of all the potential PBD-binding sites in Ana1 dramatically reduces centrosomal Polo levels

To identify proteins involved in recruiting Polo to the mother centriole we examined a small number of candidates that are important for centriole assembly and/or function in flies and that, like Polo, localise in a ring around the mother centriole: Sas-4 (CPAP), Asl (Cep152), Cep135 (also known as Bld10), Ana1 (Cep295) and PLP (Pericentrin [PCNT]) (vertebrate homologues indicated in brackets) (Mennella et al., 2012; Fu and Glover, 2012; Fu et al., 2016; Saurya et al., 2016). We previously uncovered the role of Spd-2 in recruiting Polo to the mitotic PCM by mutating all the potential S-S/T motifs in Spd-2 to T-S/T. This S-to-T substitution is conservative (French and Robson, 1983), so it is unlikely to dramatically perturb protein structure, but it abolishes PLK1 PBD-phosphopeptide binding in vitro (Elia et al., 2003a; b), and also appears to do so in vivo (Alvarez-Rodrigo et al., 2019). We generated mutant versions of all the candidate proteins in which we mutated all S-S/T motifs to T-S/T (Fig.1A). The only exception was Sas-4, for which all S-S/T motifs except the previously identified T200 motif—which has already been shown to initiate Polo recruitment at centrioles (Novak et al., 2016)—were mutated. We then analysed the centrosomal localisation of each protein and of Polo-GFP using an mRNA injection strategy (Fig.1B) (Novak et al., 2016; Alvarez-Rodrigo et al., 2019).

**Figure 1:**
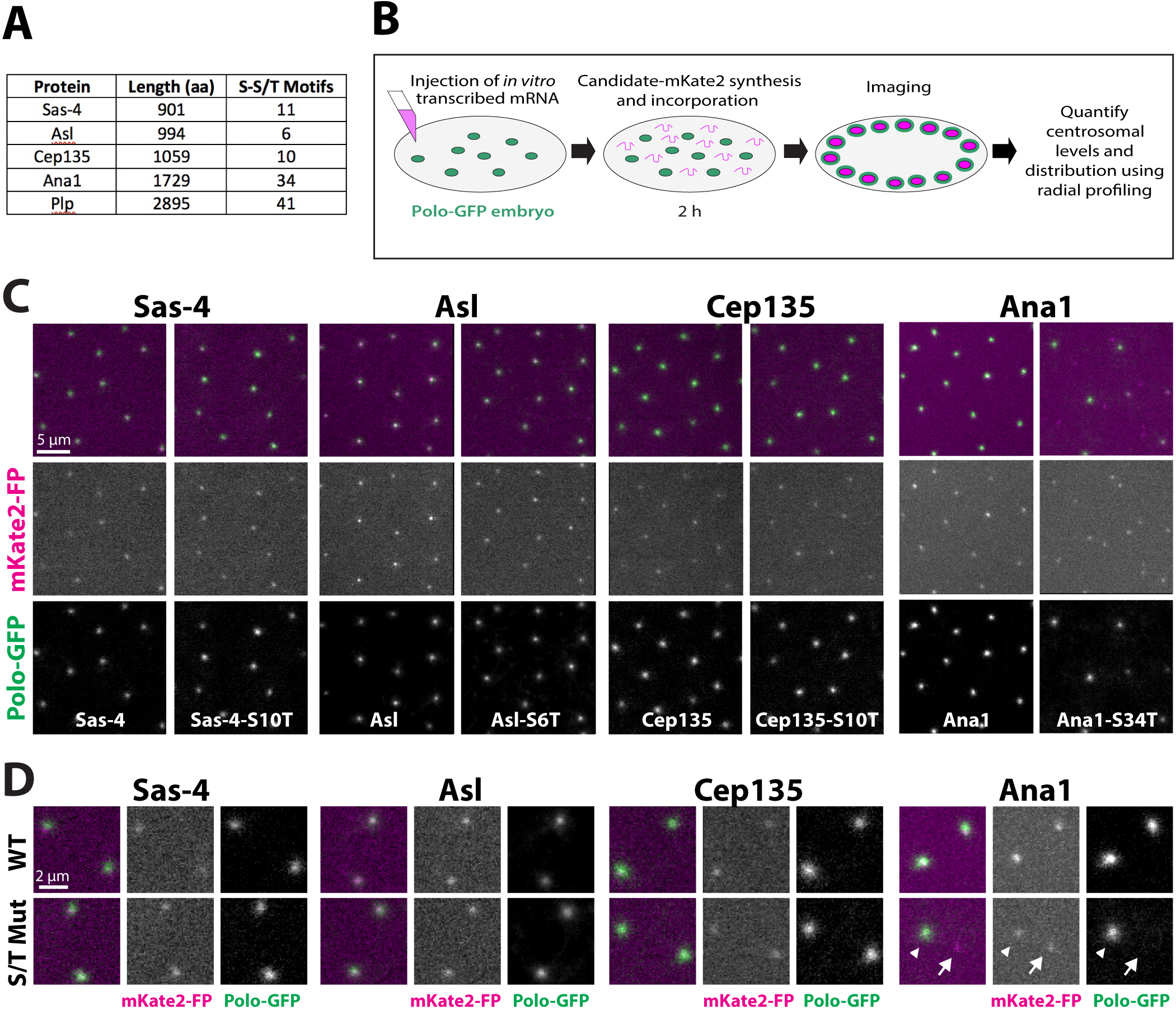
An mRNA injection-based screen for proteins that help recruit Polo to centrosomes. **(A)** Table shows the number of potential PBD-binding sites (S-S/T motifs) in several centriole proteins. **(B)** Schematic illustrates the mRNA injection assay used to test the effect on Polo recruitment of mutating all the potential PBD-binding sites in a candidate protein. *Green* circles represent centrosomes recruiting Polo-GFP. **(C)** Micrographs of embryos expressing Polo-GFP (*green* in merged channel) injected with mRNA encoding WT (left panels) or SnT mutant (right panels) forms of each of the candidate proteins (Sas-4, Asl, Cep135 or Ana1) tagged with mKate2 (*magenta* in merged channel). **(D)** Magnified view highlighting a pair of centrosomes for each condition. *Arrowheads* indicate centrosomes that contain Ana1-S34T-mKate2 and that recruit Polo-GFP; a*rrows* indicate centrosomes that contain Ana1-S34T-mKate2 but do not detectably recruit Polo-GFP. 5-14 embryos were injected and analysed for each mRNA, with >100 centrosomes visually inspected per embryos. Quantification of the recruitment of Polo-GFP in the presence of the WT and mutant Sas-4-, Asl- and Cep135-fusion is shown in Fig.S1.

We produced mRNA in vitro encoding either wild-type (WT) or the S-to-T substitution (S*n*T – with *n* indicating the number of substitutions) versions of the candidate proteins followed by a C-terminal red fluorescent-tag (mKate2), and injected this into embryos expressing Polo::GFP (Buszczak et al., 2007). The embryos were imaged 2hr after injection to allow the injected mRNA to be translated, the protein to incorporate into centrosomes, and the fluorescent tag to mature. Unfortunately, neither the WT-PLP- nor PLP-S41T-mKate2 fusion proteins were detectable at centrosomes in these experiments, perhaps because PLP is so large that more time is required for the protein to be translated and for the fluorophore to mature. PLP was therefore excluded from further analyses. All other candidate proteins were recruited to centrosomes, and the WT and S*n*T-mutant proteins exhibited qualitatively similar localisations (Fig.1C,D).

For Asl, Cep135 and Sas-4, the levels of Polo-GFP recruitment to centrosomes appeared to be similar in embryos expressing either the WT or S*n*T-mutant forms of the protein (Fig.1C,D; Fig.S1). In contrast, although Polo-GFP recruitment appeared unperturbed in embryos expressing WT-Ana1-mKate2, it was dramatically perturbed at some of the centrosomes (almost always affecting one of the two centrosomes in a separating pair) in embryos expressing Ana1-S34T-mKate2 (Fig1.C; *arrow*, Fig.1D). Due to the variability of the Polo-GFP loss at centrosomes, and the near-complete loss of Polo-GFP at many of these centrosomes, it was not possible to quantify this effect using radial profiling. Nevertheless, this striking phenotype was observed in 7/7 embryos injected with Ana1-S34T-mKate2 mRNA, compared to 0/9 embryos injected with WT-Ana1-mKate2 mRNA (scored blindly). A similar preferential loss of Polo-GFP from one centrosome in a separating pair was also observed with the Sas-4-T200 mutant protein, where it was shown that it was the new mother (**NM**) centriole that did not properly recruit Polo (Novak et al., 2016). This asymmetric behaviour is likely to occur because a fraction of the Ana1 protein, like Sas-4 (Conduit et al., 2015a), incorporates irreversibly into centrioles (Saurya et al., 2016). Thus, old mother (**OM**) centrioles can recruit more Polo presumably because they were formed when levels of the newly translated mutant protein were lower. We conclude that Ana1 helps recruit Polo to centrosomes while Asl, Cep135 and Sas-4 (apart from its previously described T200-motif, (Novak et al., 2016)) do not have a major role in this process.

### The ability of Ana1 to recruit Polo to centrioles is not required for centriole or cilia assembly

To more rigorously examine the role of Ana1 in recruiting Polo to centrosomes we generated transgenic fly lines expressing either the WT or Ana1-S34T-constructs with a C-terminal GFP- or mCherry-tag under the control of the *ubiquitin* promoter. Both the WT and mutant transgenes were equally capable of rescuing the characteristic uncoordinated phenotype of *ana1^-/-^* mutant flies— caused by the lack of centrioles/cilia in the sensory neurons (Kernan et al., 1994; Blachon et al., 2009) (Fig.2A). Moreover, *ana1^-/-^* mutant 3^rd^ instar larval neuroblasts—that normally lack detectable centrioles (Blachon et al., 2009)—exhibited normal numbers of centrioles when rescued by either the WT or Ana1-S34T-mCherry transgenes (Fig.2B). Taken together, these data indicate that despite the multiple S-T substitutions introduced into the Ana1-S34T transgene, the mutant Ana1 protein appears to support centriole duplication and cilia assembly as efficiently as the WT protein.

**Figure 2:**
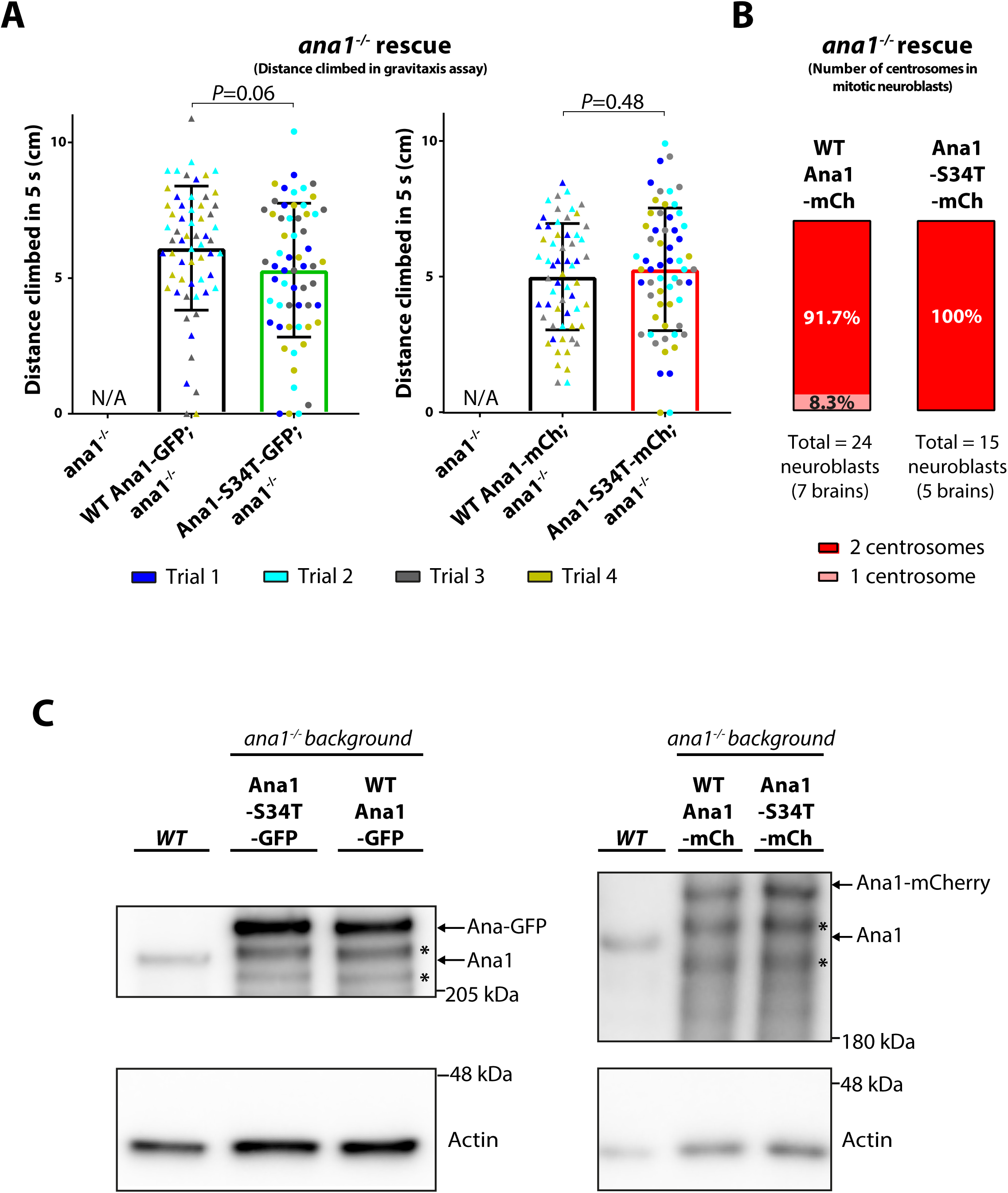
Characterisation of WT and S34T Ana1 transgenic fly lines. **(A)** Graphs show the quantification of negative gravitaxis assays (Ma and Jarman, 2011; Pratt et al., 2016). Each datapoint shows the average distance climbed by 15 individual 1–3 day-old *ana1^-/-^* male flies expressing either WT or S34T-mutant versions of Ana1—tagged with either GFP (left graphs) or mCherry (right graphs)—within the first 5 s after being tapped to the bottom of a cylinder (four technical repeats). Note that *ana1^-/-^* flies without any transgene were not scored in this assay, as 100% of the mutant flies are severely uncoordinated and so cannot climb at all. Nevertheless, we show this bar as zero—marked with not applicable (N/A)—to better illustrate the level of rescue for each transgene. Error bars represent SD. **(B)** Quantification of the percentage of mitotic neuroblasts with one or two centrosomes in *ana1^-/-^* larval brains co-expressing Spd-2-GFP and either WT Ana1-mCherry or Ana1-S34T-mCherry. Live neuroblasts were imaged and analysed blindly, with centrosomes being identified by the co-localisation of both markers. **(C)** Western blot showing total Ana1 levels in WT embryos compared to *ana1^-/-^* embryos expressing Ana1-S34T- or WT-Ana1-fusions to either GFP or mCherry driven from the *ubiquitin* promoter. Note that the anti-Ana1 antibodies recognised a ladder of proteins (*asterisks*) in the samples expressing the -GFP or -mCherry fusions that are likely to be degradation products. Actin is shown as a loading control. Each blot is representative of two technical repeats.

### Embryos expressing Ana1-S34T-transgenes die early in development

Although Ana1-S34T-fusion proteins appear to be functional for centriole and cilia assembly, *ana1^-/-^* mutant females expressing Ana1-S34T-GFP laid embryos that hatched at a frequency of only ∼0.4% (n>1000), while mutant females expressing WT-Ana1-GFP hatched at a frequency of ∼85% (n>500 embryos scored). For simplicity, **we hereafter refer to embryos laid by *ana1^-/-^* mutant females that express a WT or mutant Ana1-fusion protein as WT-Ana1 or Ana1-S34T embryos**. This was not due to the differential expression of the transgenes, as western blotting revealed that both WT and mutant Ana1-GFP or Ana1-mCherry-fusions were expressed at similar levels, and at levels that were ∼2-4X higher than the endogenous Ana1 protein in WT embryos (Fig.2C). To understand why Ana1-S34T-GFP embryos failed to hatch we fixed these embryos and stained them to reveal the distribution of microtubules (MTs), DNA, and the GFP-moiety. The nuclei in WT Ana1-GFP embryos appeared to divide synchronously, with well-formed mitotic spindles that were evenly distributed (Fig.3). In contrast, Ana1-S34T-GFP embryos displayed a range of mitotic defects that became more pronounced during the latter stages of syncytial embryo development (Fig.3). Particularly striking was the presence of many clusters of closely spaced centrosomes that appeared to be duplicated and disengaged centriole pairs that had failed to separate properly (*arrows*, Fig.3A). This interpretation was confirmed using 3D-Structured Illumination super-resolution Microscopy (3D-SIM) (see below). Most Ana1-S34T embryos became necrotic and died prior to cellularisation at nuclear cycle 14.

**Figure 3:**
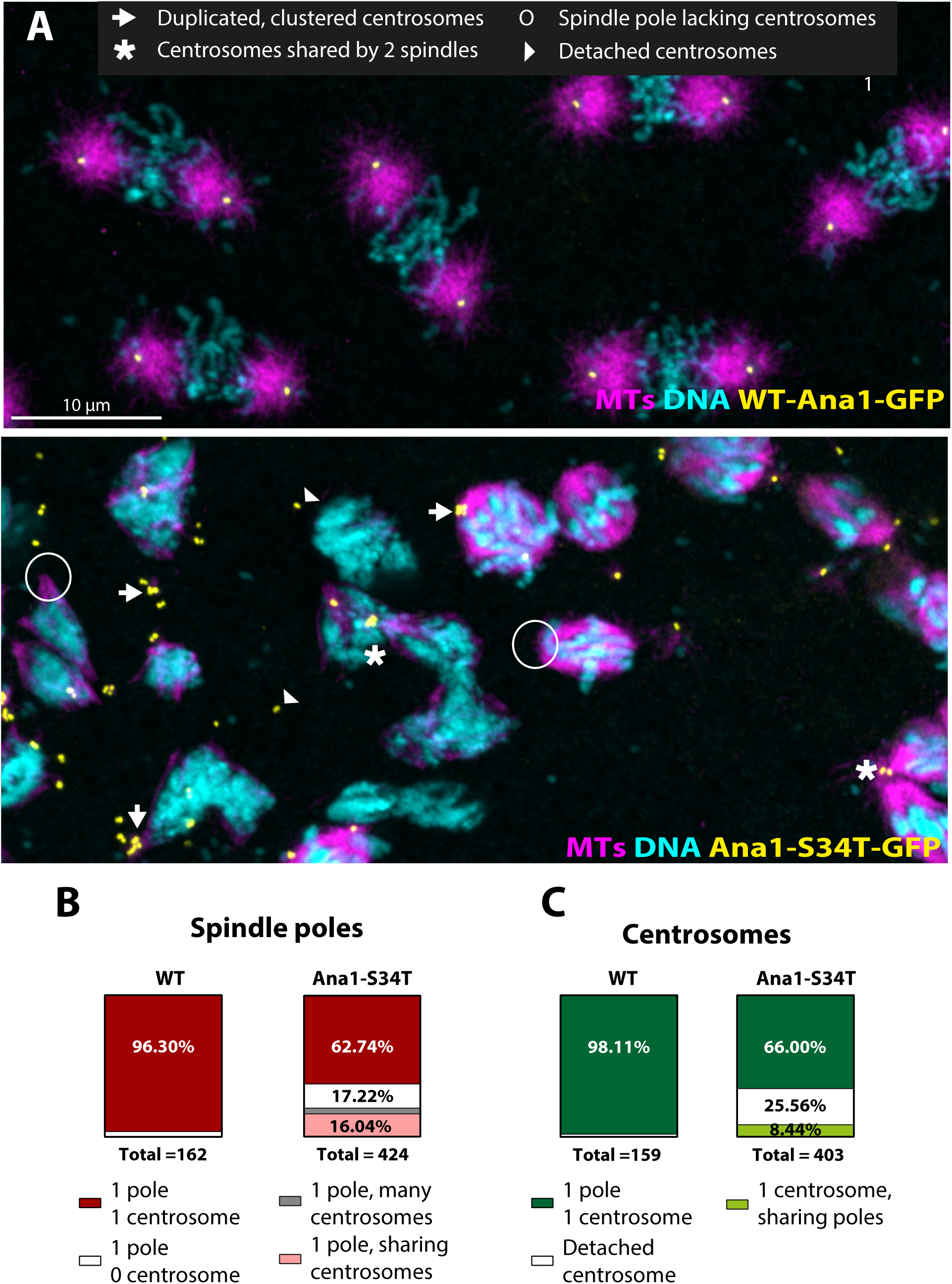
Ana1-S34T-GFP embryos accumulate severe mitotic abnormalities. **(A)** Representative micrographs of fixed *ana1^-/-^* embryos expressing WT Ana1-GFP (*top*) or Ana1-S34T-GFP (*bottom*) (*yellow*) were stained using an anti-α-tubulin antibody (to visualise MTs, *magenta*) and DAPI (to visualise nuclei, *cyan*). Mitotic abnormalities were rare in WT Ana1-GFP embryos but were frequent in Ana1-S34T-GFP embryos. Common defects observed are indicated in the figure legend and highlighted on the micrograph (only a few examples of each defect are marked to facilitate visualisation). **(B,C)** Quantification of some of the spindle pole (B) and centrosome (C) defects observed in eight WT and ten S34T-rescued embryos undergoing mitosis (one technical repeat). The total number of spindle poles and centrosomes scored per genotype is indicated.

### Ana1-S34T-GFP is recruited to centrosomes where it can recruit normal levels of Asl

We next measured the centrosomal intensity of the GFP signal in living WT-Ana1-GFP and Ana1-S34T-GFP embryos to test if the two proteins were recruited to centrioles to similar levels. WT-Ana1-GFP and Ana1-S34T-GFP both exhibited a slight asymmetry at centrosomes, being enriched in one centrosome of every pair; such asymmetry was previously reported in flies with WT-GFP-Ana1 expressed from the endogenous promoter, where it was shown that the brighter centrosome contains the OM centriole (Saurya et al., 2016). We therefore independently measured the signal intensity of the centrosomes containing the OM and NM centrioles at the beginning and end of S-phase (Fig.4A,B). Surprisingly, the centriolar levels of Ana1-S34T-GFP were always slightly higher than those of WT-Ana1-GFP. We do not understand the reason for this difference, but we conclude that the Ana1-S34T-GFP protein is recruited to centrioles just as, or even slightly more, efficiently than WT Ana1-GFP.

**Figure 4:**
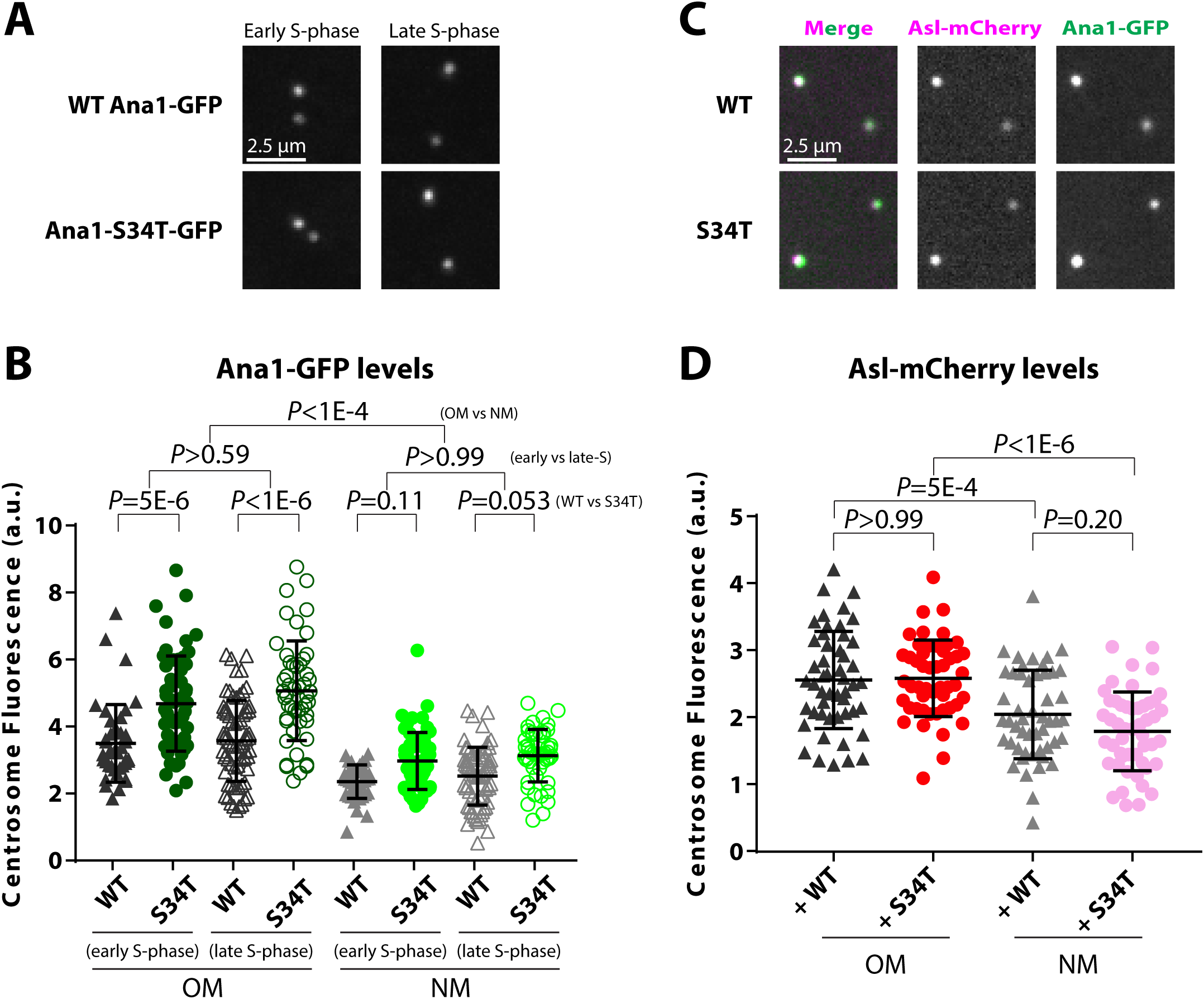
Ana1 and Asl are similarly recruited to centrioles in Ana1-GFP and Ana1-S34T-GFP embryos. **(A)** Example micrographs of WT and Ana1-S34T-GFP embryos. **(B)** Graph shows mean centrosomal Ana1-GFP intensity at the start of S-phase (*filled* symbols) and at the end of S-phase (*empty* symbols) at OM or NM centrosomes in either Ana1-GFP (*black/grey* symbols) or Ana1-S34T-GFP (*green* symbols) embryos. N = 40 (WT, early S-phase), 60 (S34T, early S-phase), 70 (WT, late S-phase) and 50 (S34T, late S-phase) centrosome pairs analysed in total per category; 10 pairs of centrosomes were analysed per embryo. For each centrosome pair, the centrosome with the highest intensity was classified as the OM (Saurya et al., 2016). **(C)** Example micrographs of WT and Ana1-S34T-GFP (*green*) embryos co-expressing Asl-mCherry (*magenta*). **(D)** Graph shows mean Asl-mCherry intensity at OM and NM centrosomes in embryos expressing either Ana1-GFP (*black/grey* symbols) or Ana1-S34T-GFP (*red/pink* symbols). Five embryos were analysed per genotype, and 10 pairs of centrosomes were analysed per embryo (so n = 50 centrosome pairs per category). For each pair, the centrosome with the highest mean Asl-mCherry intensity was classified as the OM (Novak et al., 2014). Error bars represent SD.

It has previously been shown that Ana1 helps to recruit and/or maintain Asl at mother centrioles (Fu et al., 2016; Saurya et al., 2016); this function is important, as Asl is required to allow mother centrioles to both duplicate and organise mitotic PCM (Cizmecioglu et al., 2010; Dzhindzhev et al., 2010; Hatch et al., 2010; Kim et al., 2013; Conduit et al., 2014b; Novak et al., 2014; Shimanovskaya et al., 2014). We wondered, therefore, whether the defects we observe in Ana1-S34T-GFP embryos might be due to a failure to properly recruit and/or maintain Asl. Like Ana1, Asl is preferentially concentrated at OM centrioles (Novak et al., 2014), so we quantified the levels of centriolar Asl-mCherry at OM and NM centrioles in WT-Ana1-GFP and Ana1-S34T-GFP embryos. Perhaps surprisingly, there was no significant difference between WT and S34T embryos in the levels of Asl-mCherry at the OM centrioles or between WT and S34T embryos at NM centrioles (Fig.4C,D). We conclude that the role of Ana1 in recruiting/maintaining Asl at centrioles is independent of its role in recruiting Polo to centrioles.

### Ana1 helps to recruit Polo to the mother centriole

We next tested whether centrosomal Polo recruitment was impaired in Ana1-S34T-GFP embryos, as would be expected from the results of our earlier mRNA injection experiments. We expressed Polo-GFP in embryos laid by *ana1^-/-^* mutant females that expressed either WT-Ana1-mCherry or Ana1-S34T-mCherry. The WT Ana1-mCherry embryos expressing Polo-GFP developed normally (Fig.5A), but the co-expression of Polo-GFP and Ana1-S34T-mCherry led to mitotic defects in embryos that were much more severe than those observed when mutant GFP- or mCherry-fusions were expressed in the absence of Polo-GFP, suggesting that the GFP-tagged Polo sensitises the embryos to the expression of Ana1-S34T (Fig.5B). Interestingly, we observed a similar sensitisation when Polo-GFP was co-expressed with mutant forms of Spd-2 that were unable to recruit Polo to the mitotic PCM (Alvarez-Rodrigo et al., 2019). Although *ana1^-/-^* females expressing Polo-GFP and Ana1-S34T-mCherry were not uncoordinated—suggesting that centriole duplication and cilia assembly are not perturbed in these flies—very few of the embryos they laid developed at all, and the ones that did had very few spindles or centrosomes (Fig.5B). As expected from our previous analyses, centrosomal levels of Ana1-S34T-mCherry were comparable to WT-Ana1-mCherry (Fig.5C), but centrosomal levels of Polo-GFP were dramatically reduced in embryos expressing the mutant protein (Fig.5D). Note that because we could not accurately stage the Ana1-S34T-mCherry expressing embryos, we compared their centrosomal Polo-GFP levels to a panel of embryos expressing WT-Ana1-mCherry at different nuclear cycles and at different stages of the cell cycle; centrosomal Polo-GFP levels were significantly higher (*p*-value < 10^-6^) in the WT-Ana1-mCherry expressing embryos at all stages of development and at all stages of the cell cycle (Fig.S2).

**Figure 5:**
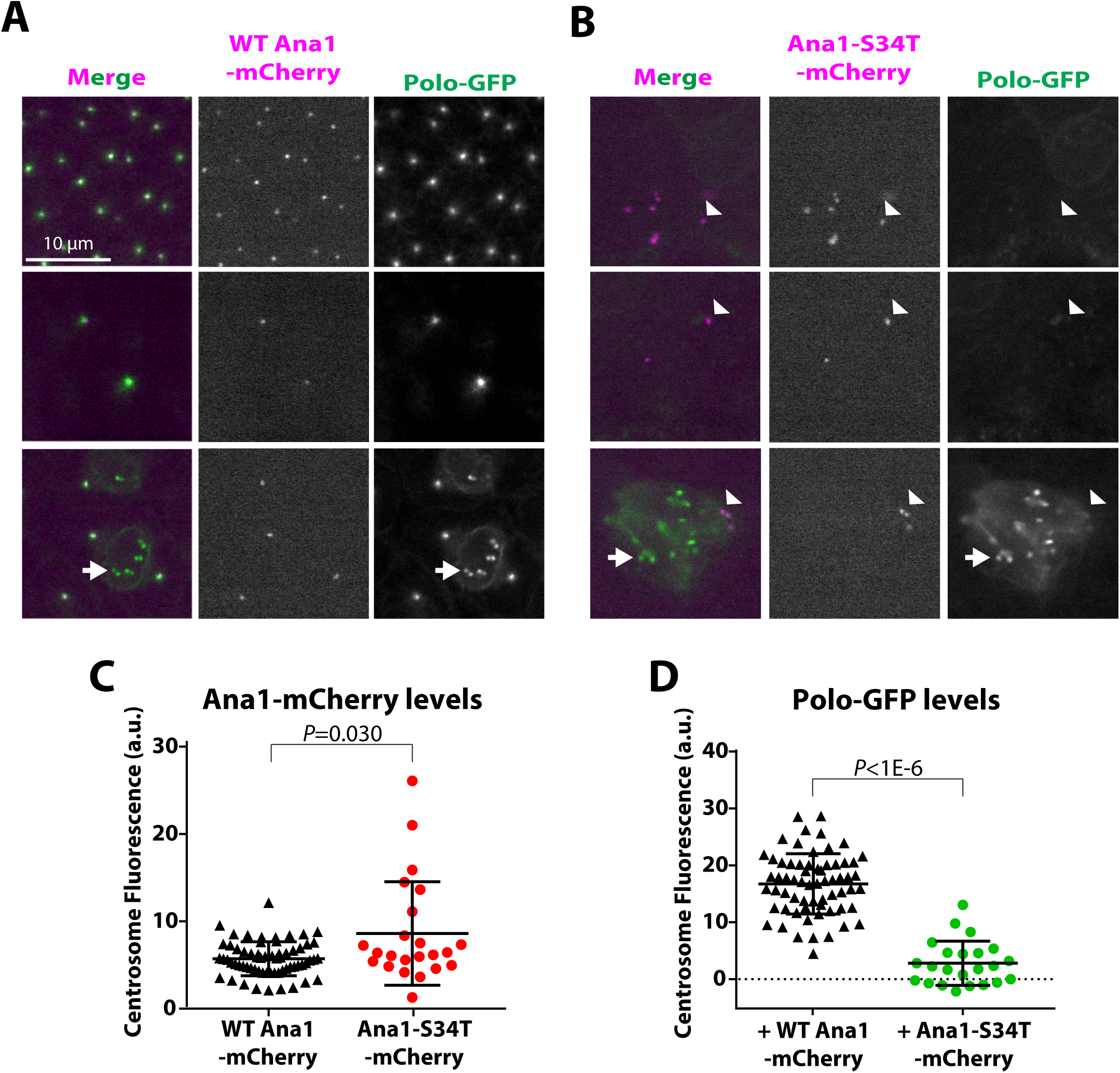
Centrosomal Polo-GFP levels are extremely low in Ana1-S34T-mCherry embryos. (**A,B**) Examples of conventional spinning disk confocal images from living WT (A) and S34T (B) Ana1-mCherry (*magenta*) embryos co-expressing Polo-GFP (*green*). Polo-GFP localised strongly to centrosomes in WT Ana1-mCherry embryos (A) but it was essentially undetectable at the centrosomes in Ana1-S34T-mCherry embryos (*arrowheads*, B)—although it was strongly recruited to other mitotic structures in the embryo (e.g. kinetochores, *arrows*). **(C,D)** Graphs show the mean centrosomal Ana1-mCherry (C) and Polo-GFP (D) intensities WT Ana1-mCherry (*black*) or Ana1-S34T-mCherry embryos (*red* and *green*, respectively). In total, n = 64 centrosomes from 10 different embryos co-expressing Polo-GFP and WT-Ana1-mCherry and 23 centrosomes from six different embryos co-expressing Polo-GFP and Ana1-S34T-mCherry. Error bars indicate SD.

We wanted to directly test whether Ana1 helps recruit Polo to the centriole wall, the mitotic PCM, or both. We could not, however, efficiently use 3D-SIM to image centrosomes in living Ana1-S34T-mCherry embryos co-expressing Polo-GFP as most embryos died too early in development. We therefore used 3D-SIM to image fixed embryos that did not express Polo-GFP. None of the commercially available anti-Polo antibodies that we tested detectably recognised Polo in fixed embryos. We therefore used an antibody that specifically recognises a phospho-epitope in Cnn (Cnn-pS567) that is phosphorylated by Polo (Feng et al., 2017) as a proxy for Polo activity at the centrosome. The presence of centrosomal Cnn-pS567 correlates well with the presence of Polo-GFP at centrosomes (Alvarez-Rodrigo et al., 2019). WT-Ana1-mCherry and Ana1-S34T-mCherry embryos were fixed and inmunostained to visualise total Cnn and Cnn-pS567. As shown previously (Feng et al., 2017), the centrosomes in WT-Ana1-mCherry embryos recruited a Cnn scaffold that tended to be enriched with Cnn-pS567 in its inner regions (i.e. the regions slightly closer to the centriole) (Fig.6A). Strikingly, in Ana1-S34T-mCherry embryos the centrosomes exhibited a marked heterogeneity: all of the centrosomes recruited at least some Cnn, but while many of the centrosomes in an embryo were devoid of Cnn-pS567 (*arrows,* Fig.6B)—consistent with a total loss of Polo from both the PCM and the centriole wall at these centrosomes—some had detectable levels of Cnn-pS567 (*arrowheads,* Fig.6B), and these centrosomes were usually associated with more total Cnn. This suggests that the Spd-2-Polo-Cnn positive feedback loop that drives the expansion of the PCM scaffold was at least partly functional at these latter centrosomes.

**Figure 6:**
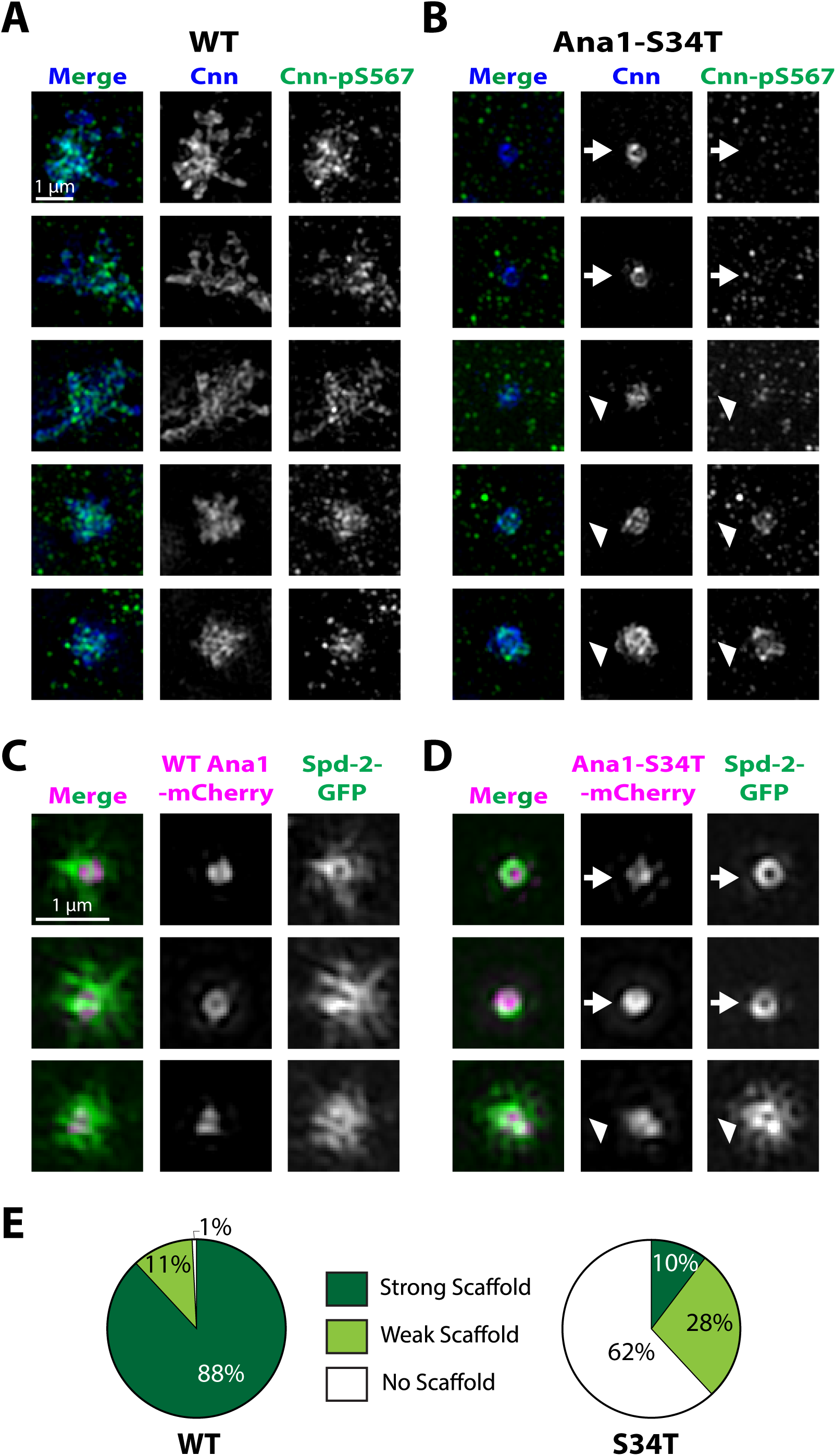
Mitotic PCM expansion is impaired in Ana1-S34T embryos. **(A,B)** 3D-SIM images from fixed WT (A) and S34T (B) Ana1-mCherry embryos. The embryos were stained with a general Cnn antibody (*blue*) and an antibody that recognises a specific Polo-dependent phospho-epitope (pSer567) in Cnn (Feng et al., 2017) (*green*). In agreement with previous reports (Feng et al., 2017; Alvarez-Rodrigo et al., 2019), most of the Cnn scaffold was phosphorylated at Ser567 in WT Ana1-mCherry embryos (A), except at the periphery of the scaffold (this was particularly noticeable during mitosis, *bottom* two panels). In Ana1-S34T-mCherry embryos (B), phosphorylated Cnn was present in some centrosomes (*arrowheads*), but not in others (*arrows*), even though unphosphorylated Cnn was present at the centriole wall in all cases. Loss of Cnn-pS567 was correlated with a lack of mitotic PCM expansion. **(C,D)** Micrographs show 3D-SIM images of individual centrosomes in living WT Ana1-mCherry (C) or Ana1-S34T-mCherry (D) (*magenta* in merged images) embryos expressing Spd-2-GFP (*green* in merged images). While Ana1-mCherry images are shown here for reference, reliably reconstructing the Ana1-mCherry signal was challenging, due to its low levels and the fast bleaching of the fluorophore. Thus, images were selected for analysis based only on whether the Spd-2-GFP reconstructed image was deemed of sufficient quality by SIM-Check (Ball et al., 2015). All centrosomes were imaged in approximately mid-S-phase when the centrosomal levels of Spd-2 are maximal. All the centrosomes in WT Ana1-mCherry embryos organised Spd-2-GFP PCM scaffolds, but this was true only in a minority of Ana1-S34T-mCherry embryos (*arrowheads*), where many centrosomes recruited Spd-2-GFP only to the centriole wall (*arrows*). **(E)** Pie charts quantify the percentage of centrosomes that qualitatively showed a strong (*dark green*), weak (*light green*) or no (*white*) pericentriolar scaffold (n = 21 for each genotype, scored blindly).

We confirmed that this was likely to be the case by examining the distribution of Spd-2-GFP in living WT-Ana1-mCherry or Ana1-S34T-mCherry embryos using 3D-SIM (Fig.6C,D). Spd-2 was concentrated at the centriole wall of all the centrosomes in these embryos, but while most of the centrosomes in the WT embryos also organised an extensive Spd-2 scaffold (Fig.6C,E), most of the centrosomes in the mutant embryos did not (*arrows*, Fig.6D; Fig.6E) although a few (∼10%) organised a relatively normal Spd-2 scaffold (*arrowheads*, Fig.6D,E). We conclude that all of the centrioles in Ana1-S34T embryos can recruit some Spd-2, and so Cnn, to the centriole wall, but only a small fraction can activate the Spd-2-Polo-Cnn positive feedback loop that drives the outward expansion of the PCM scaffold.

### Polo recruitment by Ana1 is required to efficiently activate Spd-2-Polo-Cnn scaffold assembly

We wanted to better understand the molecular basis for the puzzling heterogeneity in centrosome behaviour in Ana1-S34T embryos. We reasoned that the Polo initially recruited to centrioles by the single T200 motif in Sas-4 (Novak et al., 2016) might normally phosphorylate Ana1 to create additional PBD binding sites that serve to amplify centriolar Polo activity by recruiting additional Polo (Fig.7A). This Ana1-dependent amplification of centrosomal Polo levels might be necessary to allow Polo to phosphorylate Spd-2 to a sufficient level that Spd-2 itself can then recruit enough Polo to initiate the Spd-2-Polo-Cnn feedback loop that drives PCM expansion (Fig.7A). In such a scenario, perhaps most of the centrioles in Ana1-S34T embryos cannot recruit sufficient Polo to initiate this feedback loop, and so they fail to expand their mitotic PCM (Fig.7B[i]). Perhaps some centrioles, however, can bypass the requirement for Ana1-dependent Polo recruitment—if, for example, the centriolar Spd-2 becomes sufficiently phosphorylated (perhaps, if given enough time, by the Polo recruited by Sas-4) to initiate the feedback loop—in which case they could organise some mitotic PCM (Fig.7B[ii]). If this were the case then, once activated, the feedback loop should be self-sustaining, predicting that such “activated” centrosomes could continue to organise significant amounts of PCM scaffold through repeated rounds of division. To test this possibility, we imaged dividing centrosomes in living WT-Ana1-mCherry or Ana1-S34T-mCherry co-expressing GFP-Cnn. For these experiments we selected 8 Ana1-S34T-mCherry embryos that were at approximately nuclear cycle 11 and that were the least disrupted in their development, as this allowed us to select several centrosomes in each embryo that recruited significant amounts of PCM and that proceeded through a clear round of duplication. We compared these centrosomes to a similar set of centrosomes in similarly staged WT-Ana1-mCherry embryos.

**Figure 7:**
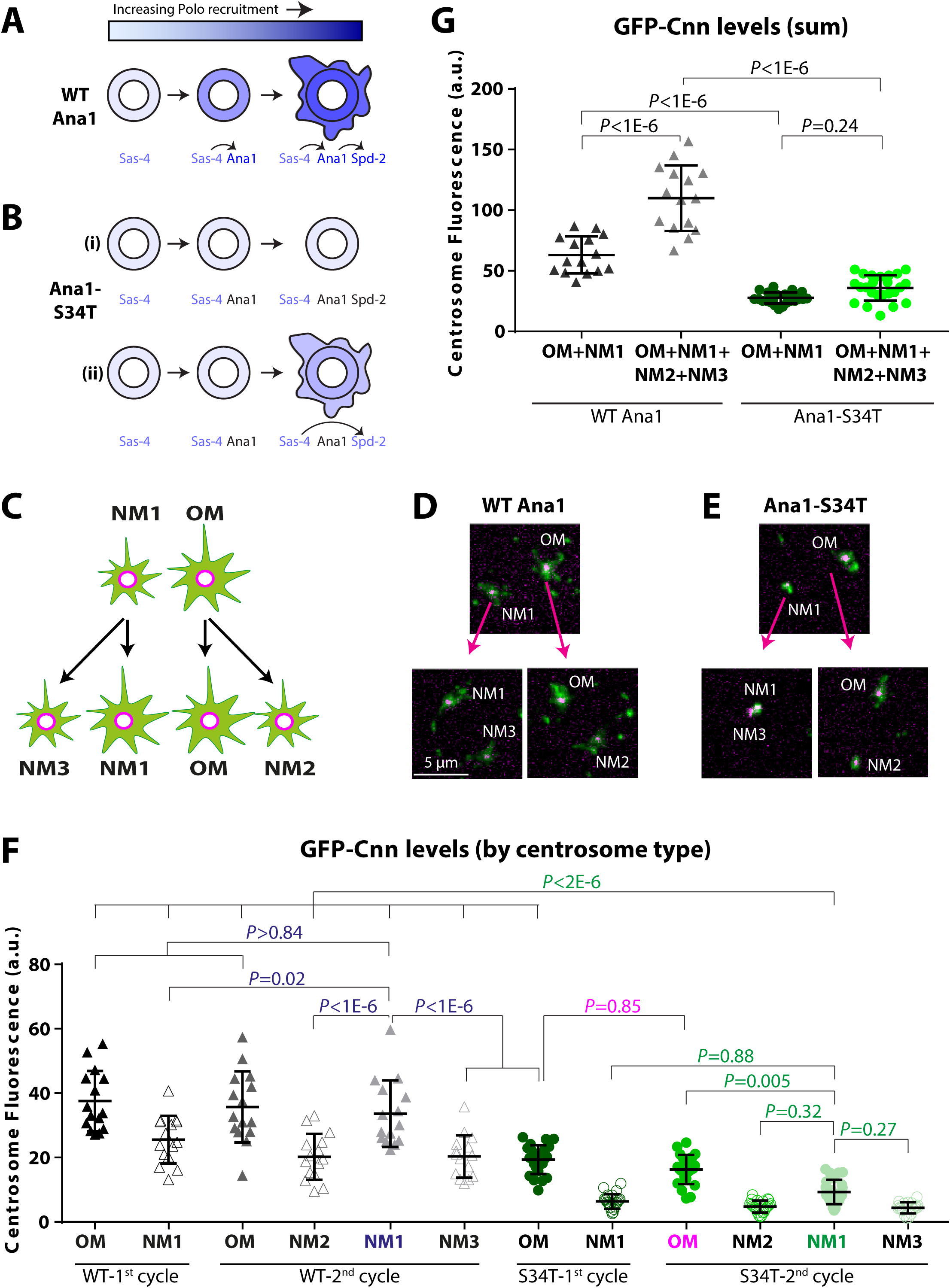
Newly formed centrosomes cannot efficiently recruit Cnn if Ana1 cannot recruit Polo. **(A,B)** Schematic illustrates how a sequential phosphorylation cascade comprising Sas-4, Ana1 and Spd-2 might drive increasing levels of Polo recruitment to the mother centriole and then to the expanding mitotic PCM in WT embryos (A), and how this process might be perturbed in embryos expressing a form of Ana1 (Ana1-S34T) that cannot efficiently recruit Polo (B). Proteins recruiting Polo are indicated in shades of blue; proteins not recruiting Polo are indicated in black. See main text for details. **(C)** Schematic illustrates the genealogy of the centrosomes analysed for their ability to recruit GFP-Cnn from one cycle to the next. In the first division cycle, the centrosome with the old mother centriole (OM) is associated with a larger GFP-Cnn scaffold than the centrosome with the new mother centriole (NM1) (Conduit et al., 2010). When these centrosomes divide, the OM and NM1 centrosomes each generate a new centrosome containing a younger mother centriole (that is again smaller than the centrosome containing the original mother centriole)—NM2 and NM3, respectively. **(D,E)** Examples of OM1 and NM1 centrosomes generated at the start of the 1^st^ cycle, and the NM2 and NM3 centrosomes they generated at the end of the 2^nd^ cycle in WT Ana1-mCherry (D) or Ana1-S34T-mCherry (E) (*magenta*) embryos expressing GFP-Cnn (*green*). In Ana1-S34T-mCherry embryos the centrosome pairs (particularly NM1+NM3) sometimes failed to separate properly. **(F)** Graph shows the mean GFP-Cnn intensity at each centrosome type in WT Ana1-mCherry (*black/grey* symbols) and Ana1-S34T-mCherry (*green* symbols) embryos. N = 5 and 8 embryos analysed, respectively; three pairs of centrosomes in the 1^st^ cycle were analysed per embryo, so a total of n = 15 and 24 centrosome pairs for each WT and S34T genotype, respectively (note that for S34T genotype, only 23 OM-NM2 pairs and 19 NM1-NM3 pairs could be analysed, due to the lack of centrosome separation at the beginning of the 2^nd^ cycle). To facilitate visualisation, only the *p-*values corresponding to the most informative statistical comparisons are shown, coloured by the type of centrosome being compared against others. **(G)** Graph shows the same data as in (F), but expressed as the average sum of GFP-Cnn levels for OM+NM1 centrosomes in the first cycle (*dark grey* for WT-rescued embryos, *dark green* for S34T-rescued embryos), and the average sum of GFP-Cnn levels for OM+NM1+NM2+NM3 centrosomes in the second cycle (*light grey* for WT-rescued embryos, *light green* for S34T-rescued embryos). Error bars represent SD.

GFP-Cnn was initially asymmetrically distributed on centrosome pairs in both WT-Ana1-mCherry and Ana1-S34T-mCherry embryos, consistent with previous reports that OM centrioles initially associate with more GFP-Cnn than NM centrioles (Conduit et al., 2010) (Fig.7C-E). We therefore refer to the larger centrosome as the OM, and the first smaller centrosome that it generates as the first NM (NM1). In WT-Ana1-mCherry embryos the OM centrosomes divided again to generate a second new mother (NM2), while the original NM1 centrosome divided again to generate a new NM centrosome (NM3). Importantly, both of the new centrosome pairs (OM-NM2 and NM1-NM3) exhibited a similar asymmetry in their GFP-Cnn levels to the original OM-NM1 pair (Fig.7F). This indicates that in these embryos all the mother centrioles recruit significant amounts of GFP-Cnn prior to their division; this increase could be quantified by comparing the sum amount of GFP-Cnn at the original centrosomes (OM+NM1) to that at the four duplicated centrosomes (OM+NM1+NM2+NM3) (Fig.7G).

In Ana1-S34T-mCherry embryos, even the OM centrosomes that we selected for their ability to recruit GFP-Cnn were associated with much less GFP-Cnn than the OM centrosomes in WT-Ana1-mCherry embryos, and the NM1 centrioles were associated with even lower levels (Fig.7E,F). Thus, even if some OM centrioles can activate the Spd-2-Polo-Cnn positive feedback loop in these mutant embryos, it is clear that they cannot do so to the normal extent. Nevertheless, these OM centrioles retained similar levels of GFP-Cnn after their division, although they again generated a NM2 centrosome that was associated with very little GFP-Cnn. In contrast, the dividing NM1 centrosomes had not grown significantly in size, and so they remained associated with very little GFP-Cnn and also generated a NM3 centrosome that was associated very little GFP-Cnn (Fig.7E,F). These data suggest that the mother centrioles in these embryos recruit very little extra GFP-Cnn prior to their division— and this could be seen by comparing the sum amount of GFP-Cnn at the original centrosomes (OM+NM1) to that at the four duplicated centrosomes (OM+NM1+NM2+NM3) (Fig.7G). We conclude that OM centrioles that organise appreciable amounts of GFP-Cnn in these embryos can continue to do so even after they have divided. We performed a similar pedigree analysis in living WT-Ana1-mCherry or Ana1-S34T-mCherry embryos co-expressing Spd-2-GFP and obtained very similar results (Fig.S3A-D).

Taken together, these experiments indicate that some OM centrosomes in Ana1-S34T-mCherry embryos can recruit significant amounts of Cnn and Spd-2, and they retain this ability even after they have divided. In contrast, most NM centrosomes cannot recruit significant amounts of PCM, at least over the timescale of our observations. These findings are consistent with our original hypothesis, and strongly suggest that some older centrosomes can bypass the requirement for Ana1-dependent Polo recruitment and activate the Spd-2-Polo-Cnn positive feedback loop that drives PCM expansion around the centriole (although at a lower level than normal) (Fig.7B[ii]).

Interestingly, we noticed that the NM1+NM3 centrosome pairs (that both failed to recruit significant amounts of Cnn or Spd-2) (Fig.7E,F; FigS3B,C) often failed to separate properly, perhaps because they collectively organised too little PCM to nucleate sufficient MTs to drive centrosome separation (Fig.7E, Fig.S3B). This presumably explains why in fixed embryos expressing Ana1-S34T-fusion proteins we observe so many centrosome clusters where mother centrioles had duplicated, but failed to properly separate from their daughters (now new mothers) (*arrows*, Fig.3A). 3D-SIM imaging of these clusters of centrioles revealed that they contained multiple disengaged Spd-2-positive centrioles, confirming that they are mothers that failed to separate properly (Fig.S3E).

### The C-terminal region of Ana1 is required to recruit Polo to centrioles

We previously found that S-S/T motifs in both the N-terminal and C-terminal region of Spd-2 contributed to recruiting Polo to the mitotic PCM (Alvarez-Rodrigo et al., 2019), so we wanted to test if this was also the case for Ana1. We therefore generated mutant forms of Ana1 which contained S- to-T substitutions in only the N-terminal (aa1-756), mid (aa756-935), or C-terminal (aa935-1729) regions (Fig.8A) and tested their ability to recruit Polo using our mRNA-injection assay (Fig.1B). Only the C-terminally mutated protein (containing 20 S-T substitutions) appeared to significantly perturb Polo-GFP recruitment, and it did so in a very similar manner to the fully mutated Ana1 protein (which contained 34 S-T substitutions) (Fig.8B).

**Figure 8:**
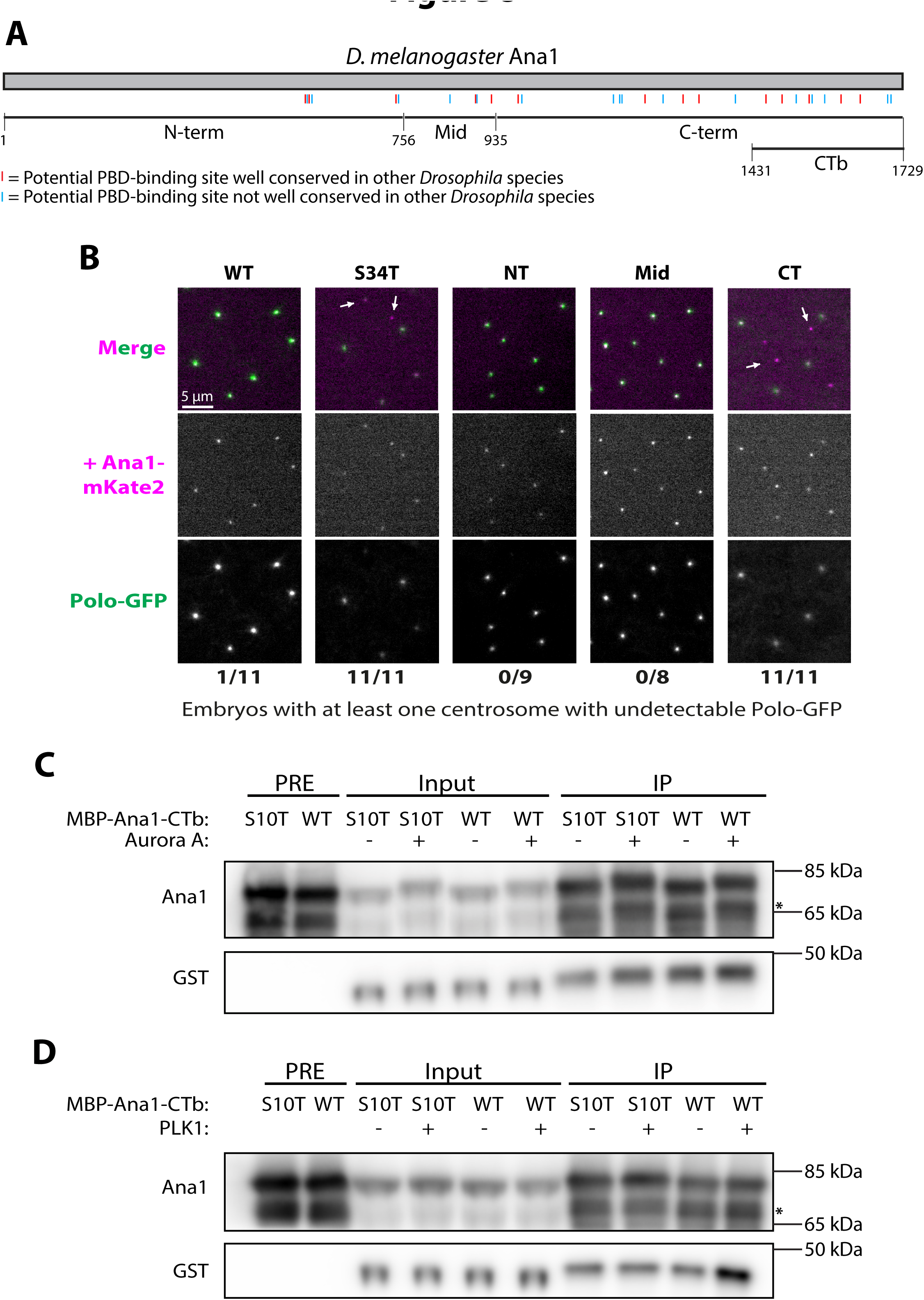
The C-terminal region of Ana1 helps recruit Polo to centrosomes and can interact directly with the PLK1-PBD. **(A)** Schematic representation of the protein sequence of *Drosophila melanogaster* Ana1 indicating S-S/T motifs that are either conserved (present in at least 11/12 *Drosophila* species analysed, *red* lines) or not conserved (*blue* lines). The boundaries of the Ana1 regions defined previously (Fu et al., 2016) and the protein fragment used for in vitro studies (CTb) are also indicated. **(B)** Micrographs of embryos expressing Polo-GFP (*green* in the merged channel) injected with constructs encoding full length Ana1-mKate2 (WT), or in which either the whole protein (S34T) or the NT, Mid or CT regions have had their S-S/T motifs replaced with T-S/T (*magenta* in the merged channel). *Arrows* highlight centrosomes that do not recruit Polo-GFP. Embryos were scored blindly for loss of Polo-GFP from at least one visible centrosome, the results for each injected construct are indicated as affected embryos/total injected embryos analysed. **(C,D)** Western blots of in vitro GST-PBD binding assays in which purified recombinant WT MBP-Ana1-CTb or MBP-Ana1-CTb-S10T (a mutant for of the CTb fragment in which all 10 S-S/T motifs have been mutated to T-S/T) were incubated with either recombinant Aurora A or PLK1 (or only kinase buffer, as a negative control) and then assessed for their ability to bind to a recombinant human GST-PBD protein. Samples were assessed prior to incubation with kinase (PRE), after incubation with the kinase and while mixed with the GST-PBD (Input), and after immunoprecipitation with anti-Ana1 antibody coated beads (IP). Both the WT and mutant MBP-fusions were not very stable and exhibited prominent degradation products (*asterisks*). Both WT and mutant MBP-fusions appeared to be phosphorylated by Aurora A (as evidenced by the shift of the proteins to slower migrating forms after incubation with the kinase) but GST-PBD exhibited a similar background binding to the WT and mutant proteins whether they had been phosphorylated or not (C). In contrast, treatment with PLK1 produced a more modest shift in protein migration, but promoted GST-PBD binding to WT MBP-Ana1-CTb but not to MBP-Ana1-CTb-S10T (D). Images are representative of two (Aurora A) and three (PLK1) technical repeats; quantification of the GST-PBD signal in all technical repeats is shown in Fig.S4.

We wanted to test whether the S-S/T motifs in this C-terminal fragment of Ana1 could interact with the PBD in vitro when phosphorylated by PLK1. We attempted to express this fragment, and several derivatives of it, as MBP-fusions in bacteria, but these were mostly insoluble and/or unstable. We did, however, manage to purify both WT and mutant forms of the very C-terminal 298aa (aa1431-1729) which contains 10 S-S/T motifs (MBP-Ana1-CTb and MBP-Ana1-CTb-S10T, respectively). We pre-treated the purified MBP-fusions with either recombinant human PLK1 or, as a control, Aurora A and then tested whether they could bind recombinant GST-PBD. Interestingly, Aurora A appeared to phosphorylate both MBP-Ana1-CTb and MBP-Ana1-CTb-S10T, as both proteins shifted to slower migrating forms after treatment with the kinase, but this did not lead to strong specific binding of GST-PBD (Fig.8C). In contrast, pre-treatment with recombinant PLK1 enhanced GST-PBD binding to MBP-Ana1-CTb but not to MBP-Ana1-CTb-S10T (Fig.8D). We remain cautious in interpreting these results as this in vitro binding assay with a small fragment of the full length protein is not representative of the situation in vivo, and it was somewhat variable (although in 3/3 repeats the overall pattern was the same and the phosphorylated MBP-Ana1-CTb protein always bound to GST-PBD most strongly). Nevertheless, in conjunction with our in vivo studies, these in vitro experiments suggest that the C-terminal region of Ana1 contains S-S/T motifs that could potentially recruit the PBD when phosphorylated in vivo, and they also indicate that PLK1 can create these recruiting sites through self-priming phosphorylation.

### The ability of Ana1 to promote centriole growth is dependent on its ability to recruit Polo

It has previously been shown that Ana1/Cep295 proteins promote centriole growth (Chang et al., 2016; Saurya et al., 2016), potentially by acting downstream of Rotatin/Ana3 to stimulate the growth of the centriole MTs as they extend distally past the central cartwheel structure (Chen et al., 2017). This “second phase” of centriole growth occurs largely in G2 and requires PLK1 (Kong et al., 2020). The centrioles in most fly cells are small and do not grow significantly during G2, but the centrioles in spermatocytes (that will go onto form the sperm flagella) elongate dramatically during an extended G2 period (Tates, 1971), and this growth requires Ana1 (Saurya et al., 2016). We therefore tested whether this growth was impaired if Ana1 cannot recruit Polo. We found that *ana1^-/-^* mutant spermatocytes expressing WT Ana1-GFP exhibited normally elongated centrioles, while those expressing Ana1-S34T-GFP were significantly shorter than normal (Figure 9A,B). These shortened centrioles, however, appeared to duplicate, disengage and separate normally so that the elongating spermatids that formed after these spermatocytes had proceeded through two rounds of meiosis appeared largely normal—although the basal bodies of the sperm flagella were significantly shorter than normal (Figure 9C,D). Motile sperm were detectable in *ana1^-/-^* mutant testes expressing Ana1-S34T-GFP, but these were qualitatively less abundant than in *ana1^-/-^* mutant testes expressing WT Ana1-GFP, and mutant males exhibited reduced fertility (data not shown). We conclude that the ability of Ana1 to promote the growth of the centrioles in spermatocytes and spermatids may be dependent on its ability to recruit Polo, but this growth is not essential for functional flagella assembly.

**Figure 9:**
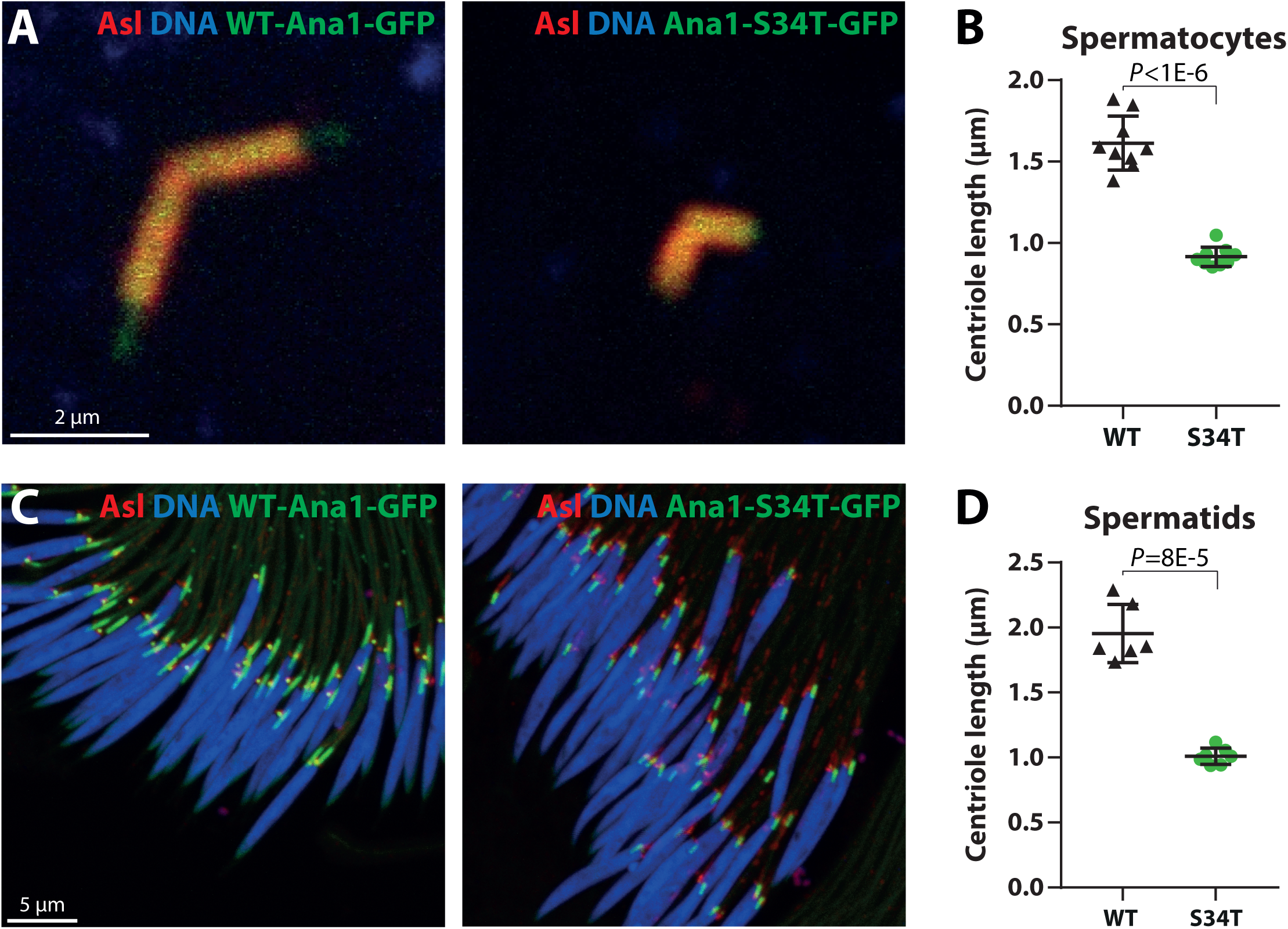
The recruitment of Polo by Ana1 is required for efficient centriole elongation in testes. **(A)** Micrographs show a typical centriole pair in *ana1^-/-^* mutant spermatocytes expressing either WT Ana1-GFP or Ana1-S34T-GFP (*green*) and stained for Asl (*red*) and DNA (*blue*). The centriole pairs in spermatocytes are engaged and arranged in a characteristic “V-shape” and they grow to a much longer size than the centrioles in most other *Drosophila* cell types during an extended G2 period. In the Ana1-S34T-GFP spermatocytes the centriole pairs are duplicated and arranged in the typical V-shape, but they are much shorter than normal. **(B)** Graphs quantify average centriole length in *ana1^-/-^* mutant spermatocytes expressing either WT Ana1-GFP or Ana1-S34T-GFP (n = 9 testes per condition, 12-30 centrioles per testis). **(C)** Micrographs show a cyst of *ana1^-/-^* mutant spermatids expressing either WT Ana1-GFP or Ana1-S34T-GFP (*green*) and stained for Asl (*red*) and DNA (*blue*). The centrioles (now extended basal bodies) are located at the distal tip of the nuclei, where they organise the sperm flagella. The mutant cysts are well organised, and the centrioles/basal bodies are correctly positioned at the base of the nuclei, but they are shorter than normal. **(D)** Graphs quantify average centriole/basal body length in *ana1^-/-^* mutant spermatids expressing either WT Ana1-GFP or Ana1-S34T-GFP (n = 6 and 7 testes respectively, 27-132 centrioles per testis). Error bars represent SD.

## Discussion

Polo has many important functions at centrioles and centrosomes and we previously showed that it is initially recruited to new-born centrioles in flies by the Cdk1-dependent phosphorylation of Sas-4 T200 during mitosis. This initial recruitment of Polo is important to allow the new-born centrioles to subsequently mature into mothers that can recruit Asl and so duplicate and recruit mitotic PCM (Novak et al., 2016). Here we show that the centriole protein Ana1 also plays an important part in recruiting Polo to mother centrioles. Our in vitro data are consistent with the possibility that Polo can phosphorylate Ana1 at one or more S-S/T motifs to “self-prime” its own recruitment. We therefore speculate that the recruitment of Polo to the maturing centriole by Sas-4 may initiate a cascade of phosphorylation events (Fig.7A). Polo recruited by Sas-4 can phosphorylate Ana1, potentially at multiple sites, to recruit more Polo to mother centrioles. The Ana1-dependent amplified pool of Polo is required to drive efficient PCM expansion, presumably because it phosphorylates Spd-2, again potentially at multiple sites. This initiates the Spd-2-Polo-Cnn positive feedback loop (Conduit et al., 2014b; Alvarez-Rodrigo et al., 2019) as phosphorylated Spd-2 can then recruit Polo to the expanding Spd-2 scaffold. Polo then phosphorylates Cnn, driving Cnn scaffold formation, the stabilisation of the Spd-2 scaffold, and the expansion of the mitotic PCM.

It is interesting to note that both Ana1/Cep295 and Spd-2/Cep192 protein families have a relatively high density of potential PBD-binding sites (S-S/T motifs) when compared to several other centriole and centrosome proteins (Fig.S5). This suggests that these proteins might have evolved to function as scaffolds that can recruit multiple PLK1 molecules to specific locations within the cell (in this case to mother centrioles and to the mitotic PCM, respectively). It will be interesting to examine whether other proteins with a high density of potential PBD-binding domains serve a similar function at other locations within the mitotic cell. Our strategy of mutating all S-S/T motifs to T-S/T in candidate proteins may be a good way of testing this possibility as, for both Ana1 and Spd-2 at least, the proteins seem to tolerate the high burden of S-to-T substitutions without significantly losing their function (apart from their ability to recruit Polo).

Importantly, our studies reveal that the ability of Ana1 to recruit Polo is also required for Ana1 to promote centriole elongation in spermatocytes during G2, but it is not required for Ana1 to help recruit and/or maintain Asl at centrioles (in contrast to the situation with Sas-4, where T200 phosphorylation is required for the proper recruitment of Asl), nor is it required for centriole disengagement. These findings indicate that either the Polo recruited by Sas-4 is sufficient for centriole disengagement, or that a separate pathway recruits Polo to centrioles to drive centriole disengagement. Moreover, Ana1 clearly has some functions that depend on its ability to recruit Polo (centrosome maturation and centriole elongation in G2), and some that do not (helping to recruit and/or maintain Asl at centrioles). Centrosome separation in G2 is also normally dependent on PLK1/Polo (Bertran et al., 2011; Mardin et al., 2011; Smith et al., 2011), and we often observed centrosomes/duplicated centriole pairs that failed to separate properly in embryos expressing Ana1-S34T. As these centriole pairs almost always organised very little PCM, however, it seems likely this that this defect is an indirect consequence of the failure to properly recruit PCM and so to nucleate the MTs that contribute to centrosome separation, rather than a direct consequence of the inability of Ana1 to recruit Polo.

These new findings strongly support our hypothesis that Spd-2, Polo and Cnn form a positive feedback loop that drives the expansion of the mitotic PCM around the mother centriole. A key feature of this model is that the Spd-2 scaffold is only recruited and assembled around the mother centriole and it then fluxes outwards: the Spd-2-scaffold itself, and the Cnn scaffold that maintains it, cannot recruit new Spd-2 into the scaffold (Conduit et al., 2014b). This is important as it can explain why the mother centriole is required to drive mitotic PCM assembly (Bobinnec et al., 1998; Basto et al., 2006; Cabral et al., 2019), and how centrioles ultimately determine mitotic PCM size (Kirkham et al., 2003; Conduit et al., 2010; Zwicker et al., 2014). Both of these findings could be explained if the mother centriole is the only source of the Spd-2 scaffold. Our observation that the pool of Polo recruited by Ana1—which, unlike Spd-2, is not a PCM component, and is restricted to the centriole— is required for the efficient expansion of the PCM demonstrates that the PCM-associated pool of Polo (recruited by Spd-2) is not sufficient to drive efficient PCM expansion on its own. We speculate that the pool of Polo associated with the mother centriole is required to constantly generate sufficient Spd-2 scaffold to support the rapid expansion of the mitotic PCM. It is important to stress, however, that so far an outward flux of Spd-2 from the centriole has only been observed in fly embryos and cells (Conduit et al., 2014b; Conduit and Raff, 2015) and it has not been observed in C. elegans embryos (Cabral et al., 2019). Clearly it will be important to test whether such a Spd-2/Cep192 flux exists in other species.

## Materials and Methods

### Fly husbandry, stocks and handling

Flies were kept at 25°C or 18°C on Drosophila culture medium (0.77% agar, 6.9% maize, 0.8% soya, 1.4% yeast, 6.9% malt, 1.9% molasses, 0.5% propionic acid, 0.03% ortho-phosphoric acid and 0.3% nipagin). The following fly lines have been previously described: Polo-GFP protein trap (Buszczak et al., 2007), eAsl-mCherry (Conduit et al., 2015a), Ubq-GFP-Cnn (Conduit et al., 2010) and Ubq-Spd-2-GFP (Dix and Raff, 2007). Embryos without endogenous ana1 (i.e. from *ana1^-/-^* mutant females) were derived from *ana1^mecB^/Df(3R)Exel7357* hemizygous mutant mothers (Blachon et al., 2009). The mCherry and GFP Ubq-Ana1 WT or S34T transgenic lines were generated by the Fly Facility in the Department of Genetics, Cambridge (UK) via random P-element insertion of the construct of choice (containing a *w^+^* gene for selection) in to a *w118* background. Stocks were kept in 8 cm x 2.5 cm plastic vials or 0.25-pint plastic bottles. *Drosophila melanogaster* Oregon-R flies were used as a WT stock for western blotting.

Embryos were collected on cranberry-raspberry juice plates (25% cranberry-raspberry juice, 2% sucrose and 1.8% agar) supplemented with fresh yeast. Standard fly handling techniques were employed (Roberts, 1998). In vivo studies were performed using 1.5-2 h-old syncytial blastoderm stage embryos. After 0-1 h collections at 25°C, embryos were aged at 25°C for 30-60 min. When injecting mRNA, embryos were collected for 20 min, injected, and imaged after 120-150 min at 21°C (but always within the syncytial blastoderm stage of development). Prior to injection or imaging, embryos were dechorionated on double-sided tape and mounted on a strip of glue onto a 35 mm glass bottom petri dish with a 14 mm micro-well (MatTek). After desiccation for 1 min (non-injection experiments) or 3 min (pre-mRNA injection) at 25°C, embryos were covered in Voltalef oil (ARKEMA). Live imaging was performed using either the spinning disk confocal or the 3D-SIM systems described below.

### Generation of Polo binding site mutants

Potential Polo binding sites in the amino acid sequence of the candidate centrosomal proteins were identified by searching for the consensus Polo binding motif S-S/T. Site conservation was assessed using FlyBase BLAST (selecting the genus *Drosophila*) and Jalview (Waterhouse et al., 2009) for protein alignment. The mutant constructs were designed in silico and synthesised externally by GENEWIZ Co. Ltd. (Suzhou, China); the WT cDNAs were obtained from Geneservice Ltd (UK). All cDNAs were cloned into a pDONR-Zeo vector and then introduced via Gateway cloning (Thermo Fisher, 11789100 and 11791100) in pRNA-mKate2CT (Novak et al., 2016) or Ubq-GFPCT and Ubq-mCherryCT (Basto et al., 2008) destination vectors as indicated. NEBuilder HiFi assembly (NEB, E2621S) was used to produce pRNA-mKate2 plasmids encoding Ana1 “partial mutants” and to introduce fragments encoding WT or mutant Ana1 amino acids 1431-1729 into a pETM44 (EMBL) vector encoding an N-terminal His6-MBP tag.

### RNA synthesis and microinjection

The mRNA injection assay has been described previously (Novak et al., 2014). In vitro RNA synthesis was performed using a T3 mMESSAGE mMACHINE kit (Thermo Fisher, AM1348) and RNA was purified using an RNeasy MinElute kit (Qiagen, 74106) according to the manufacturer’s instructions. All RNA constructs were and stored at −80 °C and injected at a concentration of 2 mg/mL.

### Behavioural assays

#### Hatching experiments

To examine the quality of the embryonic development for various fly strains generated in this study, 1-5 h collected embryos were aged for 24 h, and the percentage of the embryos that had hatched out of their chorion was calculated. At least two technical repeats per transgenic fly line (GFP- and mCherry-tagged) were performed.

#### Negative gravitaxis experiments

A standard negative gravitaxis assay was used to assess the climbing reflexes of *ana1^-/-^* mutant flies. 15 1–3-d-old adult male flies were sharply tapped to the bottom of a 10-ml cylinder, and the maximum distance climbed by individual flies within the first 5 s after tapping was measured. The distances were calculated using Fiji (ImageJ). Measurements were repeated four times (technical repeats) for each transgenic fly line (GFP- and mCherry-tagged).

### Western blot analysis

Western blotting to estimate embryonic protein levels and the results from the in vitro interaction assays was performed as described previously (Novak et al., 2014). The following primary antibodies were used: rabbit anti-Ana1 (1:500, animal #SK4818, (Stevens et al., 2009), mouse anti-Actin (1:500, Sigma, A3853) and mouse anti-GST (1:500, Thermo Fisher, MA4-004). For visualisation, we used the SuperSignal West Femto kit (Thermo Fisher, 34095) and the following HRP conjugated secondary antibodies: swine anti-rabbit (1:3000 for embryo levels, 1:10000 for in vitro assays; Dako, P0399) and sheep ECL anti-mouse (1:3000, GE Healthcare, NA931V).

### Immunofluorescence

Embryos were collected for 1 h, aged for 1 h, and processed as described (Gartenmann et al., 2020). Testes from adult male flies expressing either WT or S34T Ana1-GFP constructs in an *ana1^-/-^* background were dissected, fixed and stained, as described previously (Roque et al., 2012). Samples were mounted onto microscopy slides with high-precision glass coverslips (CellPath). The following antibodies were used: mouse anti-α-tubulin (1:1000, Sigma, DM1a), guinea pig anti-Cnn antibody (1:1000, animal #SK3516, (Lucas and Raff, 2007)), rabbit anti-Cnn pSer567 antibody (1:500, animal #30129, (Feng et al., 2017)), guinea pig anti-Asl (1:500, animal # SKC124, (Roque et al., 2012)), Alexa 647nm anti-mouse (1:500, Thermo Fisher, A21236), GFP-Booster Atto488 (1:500, Chromotek, gba488), CF405S anti-guinea pig (1:500, Biotium, 20356), Alexa 488nm anti-rabbit (1:500, Thermo Fisher, A21206) and Alexa-568 anti-guinea pig (1:500, Thermo Fisher, A11075). For quantification of mitotic defects and testes length, we used Vectashield medium with DAPI (Vector Laboratories, H-1200), whereas for the Cnn staining, we used Vectashield medium without DAPI (Vector Laboratories, H-1000).

### Imaging and image analysis

#### Spinning disk confocal microscopy

Embryos and third instar larval larval brains were imaged at 21°C on a Perkin Elmer ERS spinning disk (Volocity software version 6.3, PerkinElmer Inc.) mounted on a Zeiss Axiovert 200M microscope using a 63X/1.4-NA oil immersion objective and an Orca ER CCD camera (Hamamatsu Photonics). 488- and 561-nm lasers were used to excite GFP and mKate2/mCherry, respectively. Confocal sections of 13 slices with 0.5-μm-thick intervals were collected every 30 s. Focus was occasionally manually readjusted in between intervals.

To quantify the centrosomal levels of Polo-GFP for most candidate mRNA injection experiments, we used ImageJ to calculate an average “radial profile” of their distribution around the mother centriole as described (Alvarez-Rodrigo et al., 2019). The 5 brightest centrosomes in each embryo were analysed (the total number of centrosomes analysed for each genotype is indicated in the corresponding figure). Each individual centrosome profile was then normalised to the average peak intensity for all the centrosomes of the control (WT) condition. However, the images obtained from embryos expressing Polo-GFP injected with Ana1-S34T-mKate2 or Ana1-CT-mKate2 could not be quantified using radial profiles because these injections result in complete loss of PoloGFP from certain centrosomes. Instead, all of the images from embryos injected with these mRNAs (or their control counterparts) were analysed as follows: a two-colour z-slice with the most centrosomes in focus was selected per embryo, the images from all the different conditions tested were blinded and randomised, and one independent scorer was asked to determine for each image whether all the centrosomes visible in the red channel (i.e. those that had incorporated WT or mutant Ana1-mKate2) were also visible in the green channel (i.e. had visible amount of Polo-GFP).

For the quantification of centrosomes in neuroblasts, *ana1^-/-^* mutant larvae expressing Spd-2-GFP and Ana1-mCherry (either WT or S34T) were blinded and randomised prior to dissection, imaging and scoring. During live imaging, mitotic centrosomes in neuroblasts were identified by the presence of at least one dot of co-localised Spd-2-GFP and Ana1-mCherry and scored. 2-4 neuroblasts were scored per brain (total number of neuroblasts and brains scored are indicated in the corresponding figure).

For quantification of centrosomal protein levels in *ana1^-/-^* embryos expressing Ana1 transgenes, we measured the mean intensity within a square of fixed size centred manually on each individual centrosome, and the mean intensity of the background near each centrosome. We then calculated the average centrosome intensity and subtracted the average background intensity per embryo. The number of embryos analysed and their cell cycle stage is indicated in the corresponding figures. For embryos expressing only Ana1-GFP, we used the maximum intensity projection of the z-stack, analysed 10 pairs of centrosomes per embryo, and classified the data into two subsets: OM (data from the brightest centrosome from each pair) and NM (data from the other centrosomes). However, to analyse embryos co-expressing two different fluorescent proteins, the protocol was adapted as required (see below).

For embryos co-expressing Ana1-GFP and Asl-mCherry, or Ana1-mCherry and Polo-GFP; we could not use the maximum intensity projection of the z-stack, and instead we selected the z-slice where the most centrosomes were in focus. For embryos co-expressing Ana1-GFP and Asl-mCherry, the images were blinded prior to quantification and the data into OM and NM subsets based on Asl-mCherry intensity levels. For embryos co-expressing Polo-GFP and Ana1-mCherry, we analysed all centrosomes containing Ana1-S34T-mCherry (i.e. a total of 23 Ana1-S34T-mCherry foci from six different embryos) and quantified the mean intensity of the corresponding area in the 488 nm channel. As this was on average four centrosomes/embryo, from every WT Ana1-mCherry control embryo we analysed two random pairs of centrosomes. As it was not possible to distinguish embryo age or cell cycle stage in the S34T-rescued embryos, we analysed a panel of control embryos as diverse as possible: embryos at nuclear cycle 10 to 14, and at early S-phase, late S-phase and metaphase/anaphase. The data is shown as an average of all analysed centrosomes together (Fig.5C-D) or classifying the control centrosomes by developmental or cell cycle stage (Fig.S2).

For embryos co-expressing Ana1-mCherry and GFP-Cnn or Spd-2-GFP, we selected random pairs of separating centrosomes at the beginning of the first nuclear division imaged based solely on the Ana1-mCherry images (to avoid selection bias) and manually tracked the pairs until the following S-phase. If the tracking was successful, both pairs of centrosomes were visible and at least one of the two pairs visibly separated, that family of centrosomes was included in the analysis. Centrosomes were classified by types according to the pedigree schematic in Fig.7C, with the older mother centrosomes corresponding to those with the highest GFP-Cnn or Spd-2-GFP intensity. Three families of centrosomes were analysed per embryo, the total number of centrosomes and embryos analysed is indicated in the corresponding figure. The results were plotted as an average per type of centrosome (OM, NM1, NM2 or NM3; in the first or second cycle imaged) (Fig.7F, Fig.S3C); or as an average cumulative intensity (Fig.7G, Fig.S3D). The cumulative intensity is the sum of the intensity values for OM and NM1 centrosomes in the 1^st^ cycle analysed; or the sum of OM, NM1, NM2 and NM3 values in the 2^nd^ cycle.

#### Quantification of mitotic defects

Fixed samples were imaged using an inverted Zeiss 880 microscope fitted with an Airyscan detector (Zeiss International, Micron Oxford). The system was equipped with Plan-Apochromat 63x/1.4-NA oil lens. The laser excitation lines used were 405nm diode, 488nm argon and 633nm diode laser. Stacks of 25 slices with 0.14-μm-thick intervals were collected with pixel size (xy) of 0.035 μm, using a piezo-driven z-positioner stage. Images were Airy-processed in 3D with a strength value of “auto” (∼6). The software used to acquire images and process the images taken in super-resolution Airyscan mode was ZEN (black edition, Zeiss International). Maximum intensity projections of the images were used to count the number of centrosomes per visible pole, and the number of poles associated with each visible centrosome. One image analysed per embryo, 6-11 embryos analysed (from a panel of embryos at different points of cell cycle and syncytial stages, as it was difficult to accurately identify cell cycle/syncytial stages in Ana1-S34T-GFP embryos).

#### 3D-SIM

3D-SIM microscopy was performed and analysed as described (Conduit et al., 2014b) on an OMX V3 Blaze microscope (GE Healthcare, Micron Oxford, 29065721) with a 60x/1.42-NA oil UPlanSApo objective (Olympus); 405-, 488- and 593-nm diode lasers, and Edge 5.5 sCMOS cameras (PCO). The raw acquisition was reconstructed using softWoRx 6.1 (GE Healthcare) with a Wiener filter setting of 0.006 and channel-specific optical transfer function. Living embryos were imaged at 21°C, acquiring stacks of 6 z-slices (0.125-μm intervals). Stacks of 13 z-slices (0.125-μm intervals) were acquired from fixed samples (phospho-Cnn staining). The images shown are maximum intensity projections. The images from the different colour channels were registered with alignment parameters obtained from calibration measurements using 1 μm to 0.2 μm TetraSpeck Microspheres (Thermo Fisher) using Chromagnon alignment software (Matsuda et al., 2018). The SIM-Check plug-in in ImageJ (NIH) was used to assess the quality of the SIM reconstructions (Ball et al., 2015).

For the qualitative analysis of Spd-2 scaffold formation, centrosome images were selected based on quality of the Spd-2-GFP reconstruction as assessed by the SIM-Check plug-in and the presence of a visible, well-formed ring corresponding to the presence of Spd-2 at the mother centriole wall. Each individual centrosome image was saved as a separate file, blinded and randomised post acquisition. The entire dataset (21 individual centrosomes per condition, two conditions) was scored independently by three different researchers not involved in any aspect of the data acquisition, and an average score was calculated.

#### Testes analysis

Fixed and stained testis slides were imaged on a confocal microscope system (Fluoview FV1000; Olympus) using a 100×1.4NA oil objective and Fluoview software (Olympus). Centriole length was measured based on the Asl staining in FIJI/Image J.

### Recombinant protein expression, purification, and in vitro interaction assay

Proteins were expressed in *Escherichia coli (E. coli)* B21 strains in LB, and purified using a pre-poured amylose column containing 4 mL amylose resin (New England Biolabs, E8021L) followed by size exclusion chromatography (protein buffer: 20 mM Tris pH 8.0, 150 mM NaCl, 0.5 mM TCEP) using an AKTA pure chromatography system with a Superdex 200 10/300 GL column attached (GE Healthcare, GE17-5175-01). The in vitro interaction experiments using recombinant Ana1 fragments and commercial GST-PBD (Sigma, SRP0360) were carried out as described previously (Alvarez-Rodrigo et al., 2019) with the following modifications due to differences in specific kinase activity between PLK1 and Aurora-A: pre-incubation at 30°C with 8.8 ng/μL of commercial PLK1 kinase (ProQinase, 0183-0000-1) or equivalent volume of PLK1 storage buffer (following manufacturer’s instructions) for 90 min; or pre-incubation at 30°C with 4 ng/μL of commercial Aurora-A kinase (ProQinase, 0166-0000-1) or equivalent volume of Aurora-A storage buffer (following manufacturer’s instructions) for 30 min.

### Statistical analysis

Prism 7 (GraphPad Software) was used for all statistical analyses. The details for quantification, statistical tests, sample numbers, the measures for dispersion and exact *p-*values are described in the main text, materials and methods, or corresponding figure legends. To determine whether the data values came from a Gaussian distribution, D’Agostino–Pearson omnibus normality test was applied. To assess if the differences between means were statistically significant, we used the unpaired t-test with Welch correction (when comparing two groups) or one-way ANOVA with Tukey’s multiple comparison test (when comparing more than two groups).

## Acknowledgements

We are grateful to Zsofia Novak, Saroj Saurya and Michael Barton for technical support and advice and members of the Raff laboratory for discussions and reading the manuscript. Additional thanks to Zsofia Novak, Laura Hankins and Edward Rea for help with the blind scoring of images. 3D-SIM microscopy was performed at the Micron Oxford Advanced Bioimaging Unit, funded by a Strategic Award from the Wellcome Trust (107457). The research was funded by a Wellcome Trust Senior Investigator Award (104575 and 215523).

## Author contributions

This study was conceptualised by I.A-R and J.W.R. Investigation was done by I.A-R and A.W. Data were analysed by I.A-R, A.W. and J.W.R., who also wrote and edited the manuscript.

## Supplementary Figure Legends

**Figure S1:**
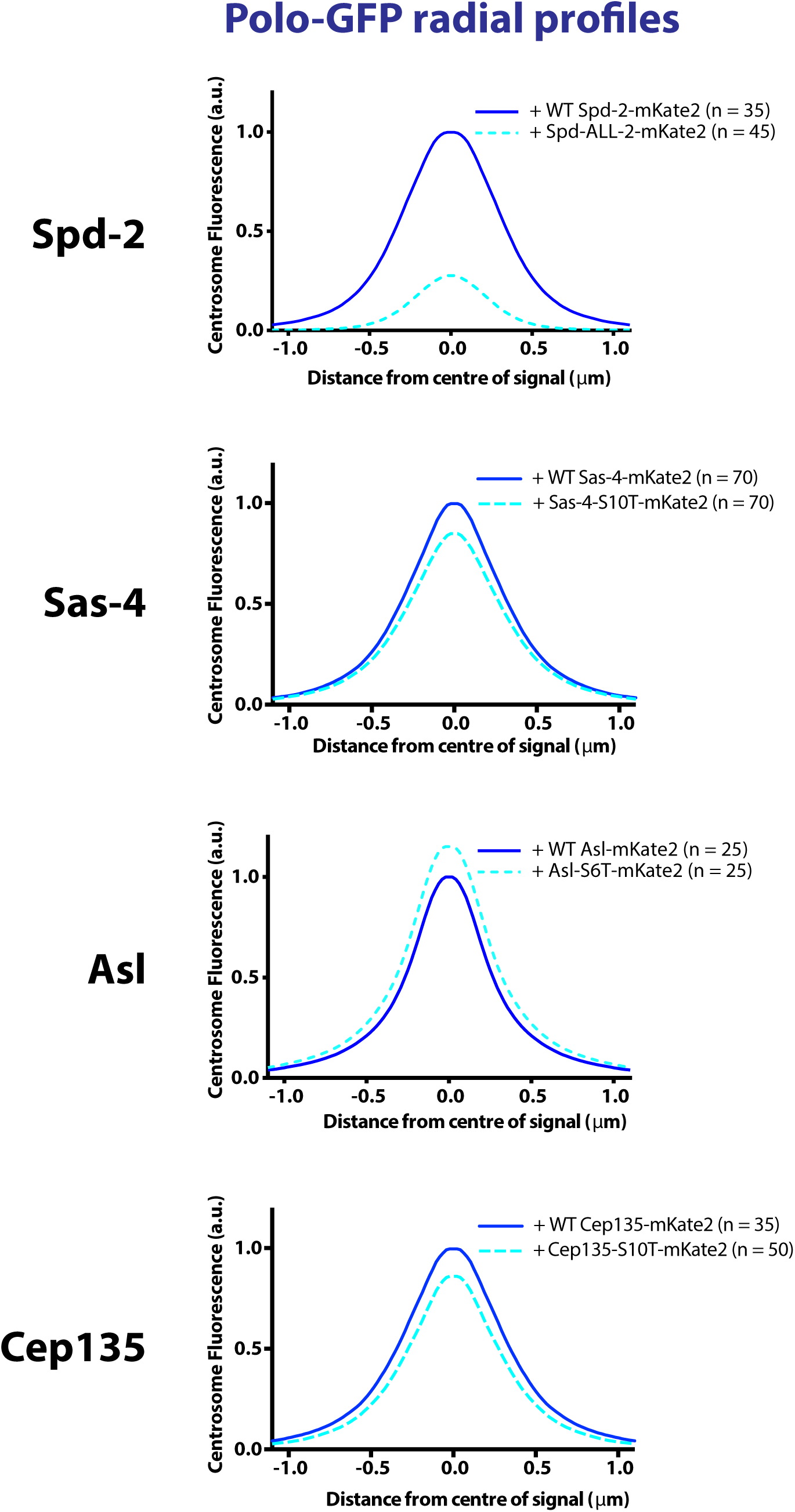
Mutating all S-S/T sites in Sas-4 (except the T200-STP motif), Asl and Cep135 has little effect on Polo-GFP recruitment to centrosomes. Radial profiles of Polo-GFP in embryos injected with mRNA encoding mKate2-tagged WT or ALL mutant Spd-2 (Alvarez-Rodrigo et al., 2019), WT or S10T Sas-4, WT or S6T Asl, and WT or S10T Cep135 (WT results are shown in *blue*, mutant results in *cyan*). The radial profiles reflect differences in protein distribution (profile width) and intensity (profile height). The effects of mutating the S-S/T motifs in Sas-4 (other than the T200-motif), Asl and Cep135 are minor compared to that of mutating all the S-S/T motifs in Spd-2 (and to the total loss observed in some centrosomes after expression of Ana1-S34T-mKate2). Five centrosomes analysed per embryo, n = number of centrosomes.

**Figure S2:**
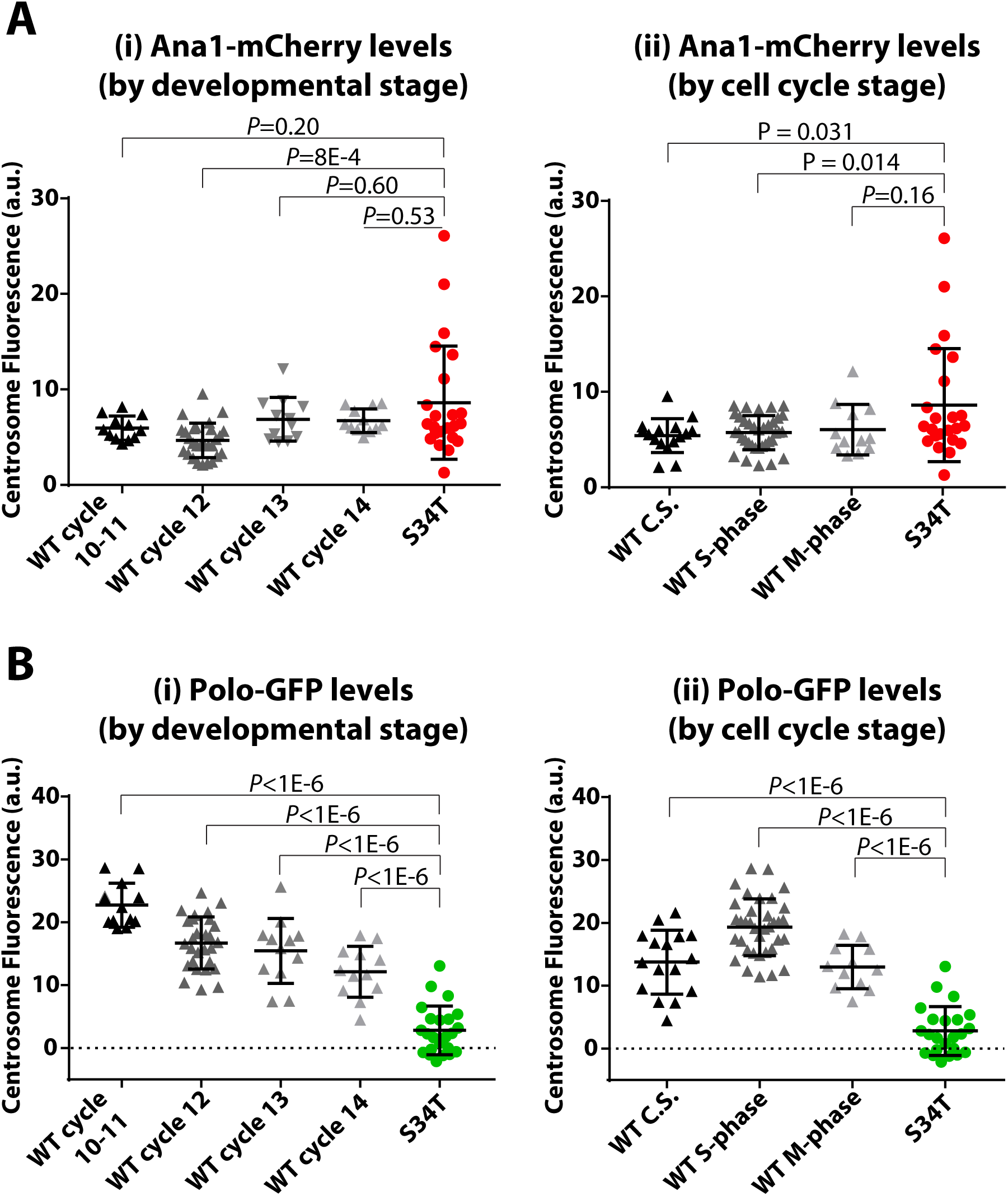
Polo-GFP levels are extremely low in Ana1-S34T-mCherry embryos compared to WT-rescued embryos at any point of the syncytial blastoderm embryonic stage or of the cell cycle. Graphs show the mean centrosomal Ana1-mCherry (A) and Polo-GFP (B) intensities in WT-rescued embryos (*dark/light grays*) and Ana1-S34T-mCherry embryos (*red* and *green*, respectively). The same data was shown in Fig.5(C,D), but here we compare the levels in S34T-mCherry centrosomes (n = 23, from six different embryos) to those from ten WT-rescued embryos classified by developmental stage (i, n = 12, 28, 12, 12) or cell cycle stage (ii, n = 16, 36, 12) (note that not all WT-rescued embryos were analysed at every cell cycle/developmental stage). C.S. = centrosome separation. Error bars indicate SD.

**Figure S3:**
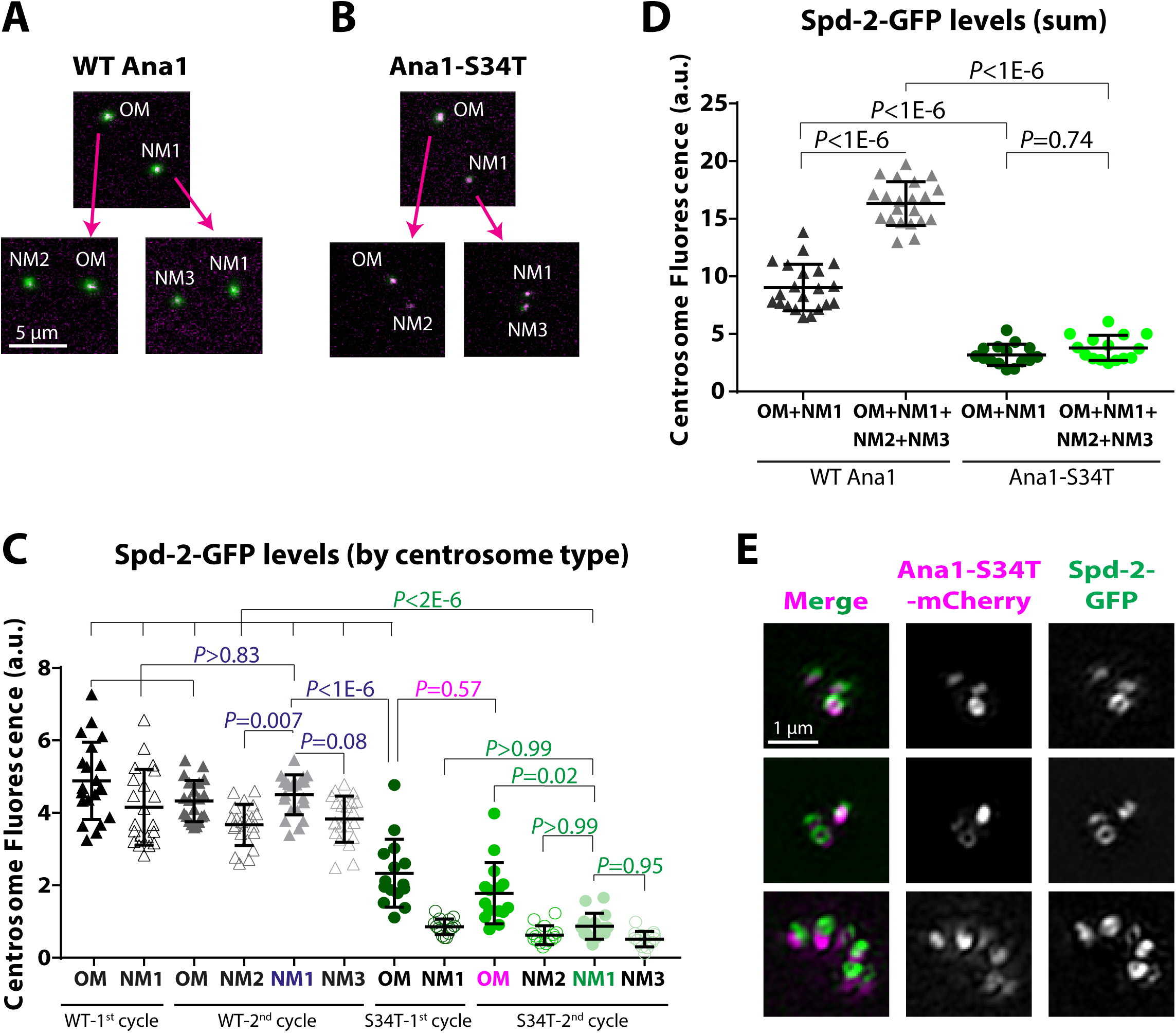
Newly formed centrosomes are unable to recruit a Spd-2 PCM scaffold if Ana1 cannot recruit Polo. **(A,B)** Examples of conventional spinning disk confocal images used for the quantification in (C,D), showing a pair of centrosomes (OM and NM1) from embryos expressing Spd-2-GFP (*green*) and WT Ana1-mCherry (A) or Ana1-S34T-mCherry (B) (*magenta*) in the first cell cycle analysed, and the four types of centrosomes that result in the second cycle analysed: OM+NM2 and NM1+NM3. **(C)** Graph showing mean Spd-2-GFP intensity above background per centrosome type for Ana1-S34T-mCherry and WT-rescued embryos (five and seven embryos analysed, respectively). Three pairs of centrosomes in the first cycle were analysed per embryo, so n = 21 for each WT centrosome type and 15 for each S34T type. As in Fig.7F, to facilitate visualisation, only the *p-*values corresponding to the most informative statistical comparisons are shown in the figure. **(D)** Graph showing the same data as (C), but expressed as the average sum of Spd-2-GFP levels for OM+NM1 centrosomes in the first cycle (*dark grey* for WT-rescued embryos, *dark green* for S34T-rescued embryos), and the average sum of Spd-2-GFP levels for OM+NM1+NM2+NM3 centrosomes in the second cycle (*light grey* for WT-rescued embryos, *light green* for S34T-rescued embryos). **(E)** Micrographs from live 3D-SIM images of centrosome clusters from *ana1^-/-^* embryos expressing Ana1-S34T-mCherry (*magenta* in merge images) and Spd-2-GFP (*green* in merge images). Ana1-mCherry images are shown for reference, the images were selected based only on whether the Spd-2-GFP reconstructed image was deemed of sufficient quality by SIM-Check (Ball et al., 2015) (see also Fig.6C,D and Materials and Methods). Spd-2 is a mother centriole specific marker, so its presence in all the centrosomes that make the clusters indicate that these are formed by mother centrioles that failed to separate properly, rather than daughter centrioles that failed to disengage from their mothers and/or disengage. Error bars represent SD.

**Figure S4:**
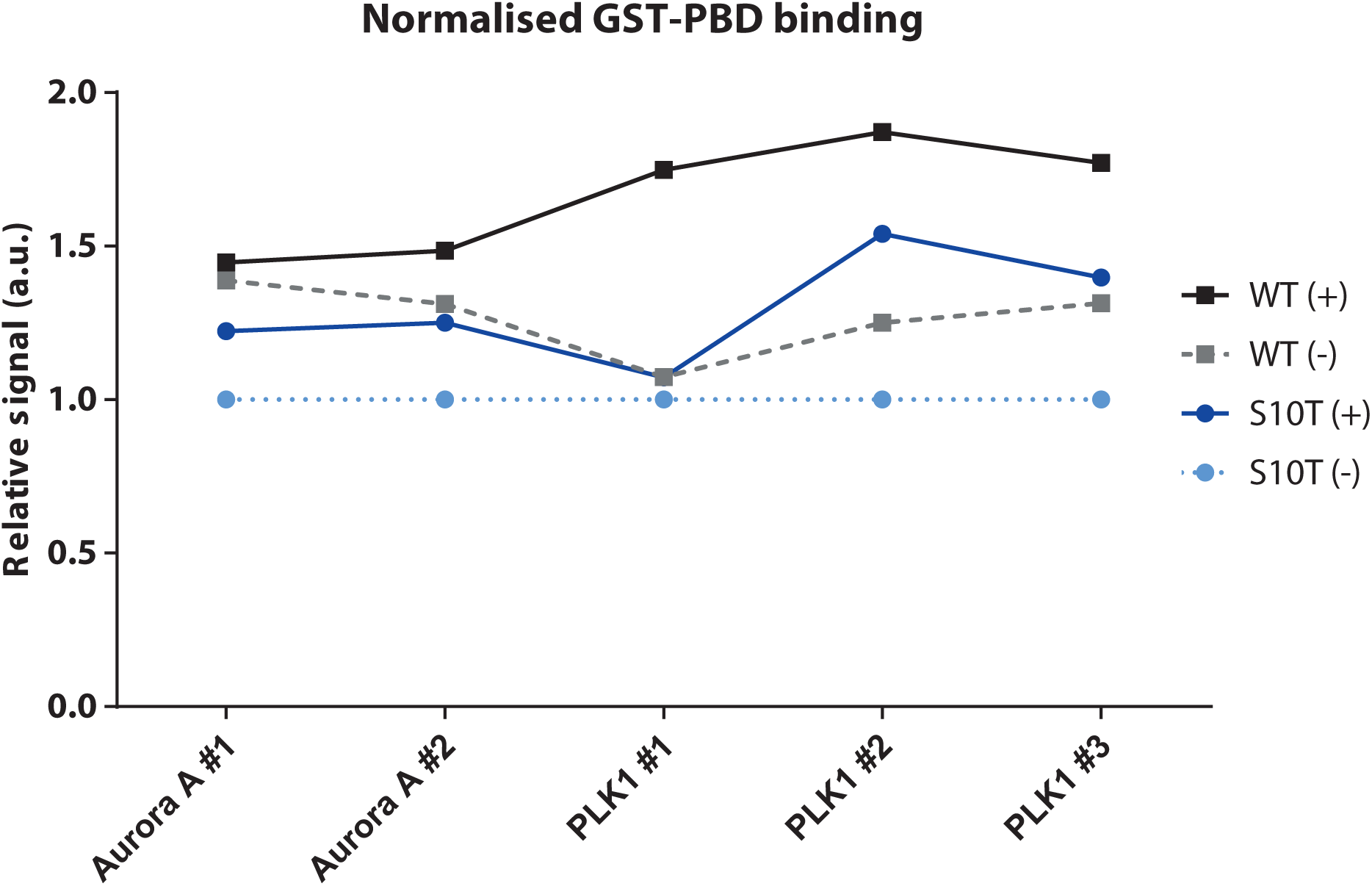
GST-PBD binds preferentially to WT Ana1 C-terminal fragments upon phosphorylation by PLK1. Quantification of the level of GST-PBD enrichment in each sample from the in vitro interaction assays, indicating the commercial kinase used for phosphorylation and the technical repeat number. The amount of GST-PBD isolated from each sample was normalised to the amount of Ana1 present in the fraction and expressed relative to the negative control (unphosphorylated MBP-Ana1-CTb-S10T). Data from the same type of sample (WT or S10T, phosphorylated or not) are shown connected to facilitate trend visualisation. GST-PBD bound preferentially to WT MBP-Ana1-CTb, particularly when the Ana1 fragment had been pre-treated with PLK1. However, due to the experimental variation and the small sample size, we were unable to test if the difference was statistically significant.

**Figure S5:**
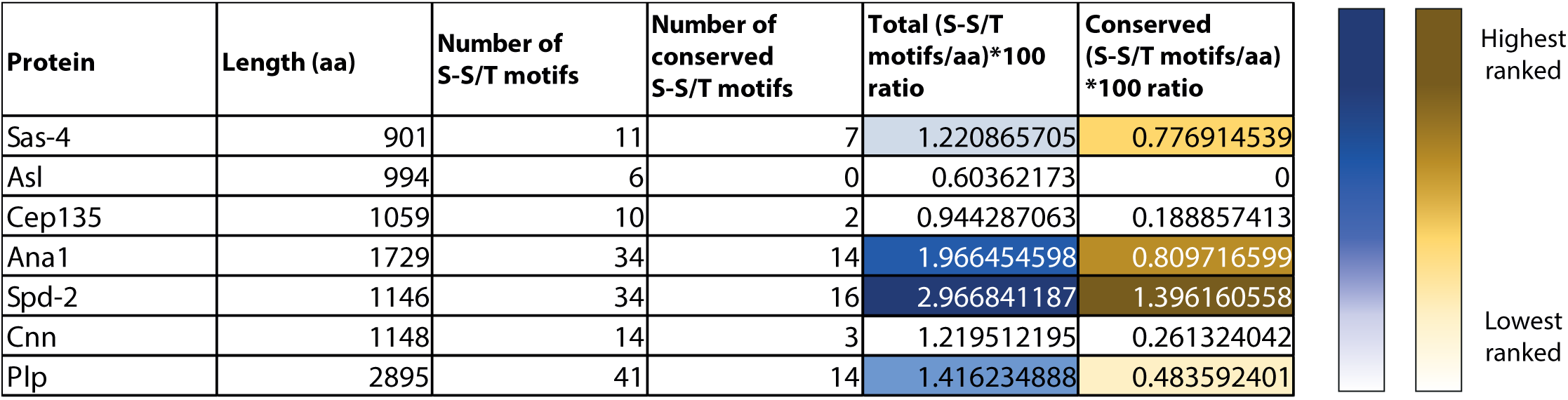
Ana1 and Spd-2 have a relatively high density of potential PBD-binding sites. Table comparing several different centrosomal proteins on the basis of their protein sequence length (expressed as number of amino acids, aa), total number of potential Polo binding sites (i.e. any S-S/T motif) in their sequence, and number of S-S/T motifs which are conserved amongst at least 11 of 12 *Drosophila* species included in this analysis. The number of total or conserved S-S/T motifs was divided by the total number of amino acids of the protein and multiplied by 100 to calculate the ratio of S-S/T motifs per 100 aa (*blue* column) and the ratio of conserved S-S/T motifs per 100 aa (*yellow* column). The highest four ratios for each column are highlighted as per the colour schemes to the right of the table (darkest tone indicating the highest ratio): Spd-2 and Ana1 (the positive hit from our assay) have the highest total and conserved S-S/T motifs/aa ratios, followed by Plp (which could not be tested using this assay) and Sas-4 (which has one site that can bind Polo).

